# PhosSight: a Unified Deep Learning Framework Boosting and Accelerating Phosphoproteome Identification to Enable Biological Discoveries

**DOI:** 10.64898/2026.03.08.708781

**Authors:** Ben Wang, Zhiyuan Cheng, Chengying She, Jiahui Zhang, Lin Lv, Hongwen Zhu, Lizhuang Liu, Yan Fu, Xinpei Yi

**Author notes:** Correspondence (L.L.); (Y.F.); (X.Y.). These authors contributed equally (Ben Wang, Zhiyuan Cheng, Chengying She).

## Abstract

Protein phosphorylation is a key regulator of signaling, with mass spectrometry (MS) based phosphoproteomics serving as the premier technology for its analysis. However, phosphorylation profiling is hindered by acquisition biases: Data-Dependent Acquisition (DDA) suffers from stochastic undersampling and missing values, while Data-Independent Acquisition (DIA) faces computational bottlenecks and inefficiencies from vast spectral libraries. We present PhosSight, a unified deep learning framework designed to augment identification depth and accelerate search efficiency. PhosSight features PhosDetect, a model that explicitly encodes phosphorylation specific physicochemical features to accurately predict peptide detectability. For DDA, PhosSight leverages predicted retention time, fragment intensity, and detectability to refine site localization and rescoring, recovering marginal, low-abundance spectra. For DIA, PhosSight utilizes detectability-guided library pruning to remove non-detectable noise, accelerating search speeds without compromising sensitivity. Benchmarking on synthetic and real world datasets confirms PhosSight’s superior performance in both modes. Applying PhosSight to a large-scale Uterine Corpus Endometrial Carcinoma (UCEC) cohort significantly reduced missing values and expanded the quantifiable phosphoproteome. This enhanced completeness enabled the discovery of novel prognosis associated kinase targets, such as MARK2, underscoring PhosSight as a powerful tool for biological discovery in precision oncology.

## Introduction

Phosphorylation is one of the most ubiquitous and functionally critical post-translational modifications (PTMs), orchestrating signal transduction, cell-cycle control, apoptosis, and metabolic reprogramming [1–3]. Its dysregulation serves as a hallmark of cancer and other pathologies, rendering the large-scale, site-specific mapping of the phosphoproteome a central objective in biomedical research [4,5]. Liquid chromatography-tandem mass spectrometry (LC-MS/MS)-based shotgun phosphoproteomics has emerged as the gold standard for achieving such global analysis [6,7]. Current acquisition strategies are broadly categorized based on precursor selection mechanisms: Data-Dependent Acquisition (DDA) and Data-Independent Acquisition (DIA) [8]. In DDA workflows, the most abundant precursor ions are sequentially selected for fragmentation [9]. While well-established, this stochastic sampling approach often leads to poor reproducibility and limited sensitivity for low-abundance phosphopeptides, resulting in significant missing values across cohorts [10]. Conversely, DIA systematically fragments all precursors within predefined isolation windows. While this theoretical comprehensiveness offers superior reproducibility and quantitative accuracy, it generates complex, highly chimeric spectra and massive datasets, which become a significant computational bottlenecks for data analysis [11].

In DDA workflows, conventional database-search strategies remain the cornerstone of identification, exemplified by widely adopted engines such as SEQUEST [12], Mascot [13], X!Tandem [14], pFind [15], MaxQuant [16], and Comet[17]. To address the specific challenge of defining exact modification coordinates, a suite of site-localization algorithms, including AScore [18], MD-Score [19], SLIP [20], PhosphoRS [21], PTM-Score [22], and PTMiner [23] has been developed. These tools employ diverse probabilistic or combinatorial principles to refine the localization confidence of phosphosites, establishing a robust baseline for DDA analysis.

Parallelly, the evolution of DIA analysis has been driven by sophisticated software suites like Spectronaut [24], DIA-NN [25], and MaxDIA [11]. These platforms increasingly rely on comprehensive spectral libraries and advanced scoring algorithms to deconvolute complex chimeric spectra. For instance, Spectronaut has become widely applicable across DIA studies due to its flexible library construction and robust data-processing capabilities, while MaxDIA offers a comprehensive workflow ranging from spectrum matching to quantitative analysis. Notably, DIA-NN has pioneered the integration of deep neural networks and novel quantification stategies to enable efficient, high-throughput data analysis. Collectively, these software tools and algorithms provide powerful support for phosphopeptide identification in DIA workflows.

Despite these methodological advances, phosphoproteomics still faces distinct bottlenecks depending on the acquisition. In DDA workflows, the primary limitation lies in identification sensitivity and depth. The labile phosphoester bond often undergoes neutral loss during fragmentation, dominating the spectra and suppressing the generation of sequence-determining ions [26]. Consequently, conventional search engines struggle to distinguish true phosphopeptides from background noise or accurately localize phosphorylation sites, leading to a high rate of false negatives and “missing” data [27]. In contrast, for DIA workflows, the bottleneck has shifted towards computational efficiency and search specificity. To achieve comprehensive coverage, DIA data must be queried against vast spectral libraries containing all theoretically possible phosphopeptidoforms. This combinatorial explosion creates an excessive search space, which not only dramatically slows down data processing but also challenges false discovery rate (FDR) control due to the increased probability of random matches. Thus, while DDA demands enhanced scoring power to rescue marginal spectra, DIA urgently requires search space optimization to handle the deluge of data.

To surmount these barriers, computational proteomics has increasingly pivoted toward deep learning. In DDA workflows, specialized models have been successfully deployed to refine identification. DeepFLR [28] utilizes predicted fragment ion intensities to estimate accurate False Localization Rate (FLR) for phosphosites. DeepRescore2 [27] integrates both fragment ion intensity and retention time (RT) predictions to enhance phosphosite localization and improve peptide rescoring. In DIA workflows, the computational challenge of expansive search spaces for unmodified peptides has been addressed by detectability models such as DeepDetect [29], which estimate peptide “detectability” to construct compact, optimized sub-libraries, thereby accelerating search speeds. Yet, a critical blind spot remains that no such detectability model currently exists for phosphorylated peptides. The addition of a phosphate group drastically alters physicochemical properties, such as ionization efficiency and solvation energy [7], rendering generic models trained on unmodified peptides (like DeepDetect) ineffective. Consequently, phosphoproteomics workflows remain unable to effectively build optimized sub-libraries or leverage detectability features for rescoring, thereby leading to inflated computational costs for DIA and suboptimal identification sensitivity for DDA.

To bridge this critical gap, we present PhosSight, a unified deep learning framework powered by our novel model PhosDetect. Unlike previous generic predictors, PhosDetect explicitly encodes phosphorylation-specific physicochemical features, improving prediction precision by up to 2.75-fold compared to existing models and establishing itself as the first accurate “detectability” predictor for the phosphoproteome. We integrated this predictive engine into PhosSight to implement a dual-mode optimization strategy. In DDA workflows, PhosSight advances the DeepRescore2 paradigm by incorporating the detectability score as a decisive, orthogonal feature. Validated on synthetic ground-truth datasets and multiple real-world cohorts, this approach boosts identification sensitivity by up to 19.5% in synthetic datasets and by over 30% in complex phosphoproteomic datasets, effectively recovering phosphopeptides missed by conventional algorithms. Furthermore, it yields an additional 8–15% gain even when compared against DeepRescore2 workflows, effectively reclaiming low-abundance signals previously lost to analytical noise. In DIA workflows, PhosSight employs PhosDetect to perform intelligent spectral library pruning. By filtering out non-detectable peptide prior to searching, the framework accelerates database search times by 40% while ensuring FDR control without compromising identification depth. Crucially, applying PhosSight to a large-scale UCEC cohort expanded the quantifiable phosphoproteome by 17% and uncovered novel prognosis-associated targets, such as PARP1_T368 and MARK2, providing new avenues for precision oncology.

## Results

### PhosSight Workflow

To address the distinct challenges of DDA and DIA phosphoproteomics, we developed PhosSight, a unified deep learning framework powered by PhosDetect, our novel phosphorylation-aware detectability predictor. PhosSight seamlessly integrates detectability scores into existing workflows to enhance identification confidence and search efficiency (**Figure 1**).

**Figure 1.**
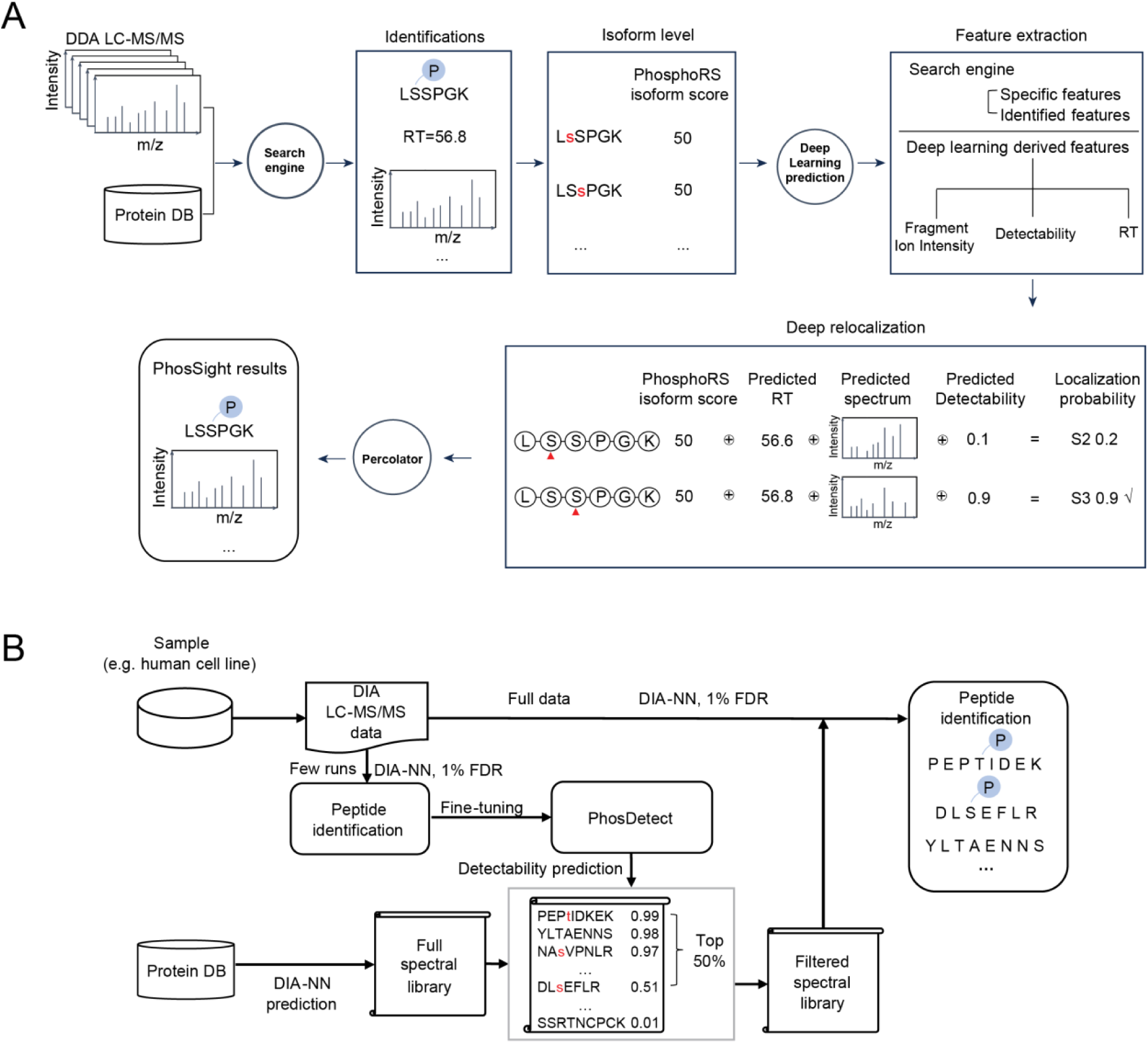
PhosSight integrates detectability scores into existing workflows to enhance identification confidence and search efficiency. A. Application in the DDA module. PhosSight operates as a post-search refinement engine for Data-Dependent Acquisition (DDA). Following initial database searching and site-localization scoring (e.g., via PhosphoRS), PhosSight extracts a multi-dimensional feature set for each peptide-spectrum match (PSM). This set integrates traditional search engine metrics with three deep-learning-derived dimensions: fragment ion intensity (predicted by pDeep3), retention time (RT, predicted by AutoRT), and peptide detectability (predicted by PhosDetect). These features are fed into a deep relocalization workflow, where detectability scores serve as critical discriminators to resolve positional isomers (e.g., distinguishing phosphorylation at S2 vs. S3) when spectral evidence is ambiguous. The refined features are finally processed by Percolator for semi-supervised rescoring to achieve high-confidence phosphopeptide identification. B. Application in the DIA module. In Data-Independent Acquisition (DIA) workflows, PhosSight implements an intelligent spectral library pruning strategy. The process begins by fine-tuning the PhosDetect model using a small subset of experimental data to capture instrument-specific ionization patterns via transfer learning. Simultaneously, a comprehensive in silico spectral library is generated from a protein database. PhosDetect then predicts the detectability of all library entries, allowing PhosSight to filter out “non-detectability” precursors (e.g., retaining only the top 50% of detectable peptides). This optimized, filtered spectral library is then used to query the full DIA dataset, significantly reducing the computational search space and accelerating processing speeds without compromising sensitivity.

In the DDA module (**Figure 1A**), PhosSight operates as a post-search rescoring and relocalization engine. First, experimental spectra are searched using standard engines (e.g., Comet, MaxQuant) combined with PhosphoRS for initial site localization (**Methods**). PhosSight then extracts a comprehensive feature set (**Supplementary Table 1**) for each peptide-spectrum match (PSM), integrating three deep learning-derived dimensions: (1) fragment ion intensity predicted by pDeep3 [30], (2) retention time (RT) predicted by AutoRT [31], and crucially, (3) peptide detectability predicted by PhosDetect. These features drive a deep relocalization process. As illustrated in Figure 1A, for ambiguous phosphopeptidoforms (e.g., phosphorylation at S2 vs. S3) where spectral evidence is marginal, the detectability score serves as a critical discriminative feature, prioritizing the peptidoforms with higher physicochemical stability and ionization potential (e.g., S3 with a score of 0.9 vs. S2 with 0.1). Finally, all features are fed into Percolator for semi-supervised rescoring (**Methods**), yielding a high-confidence dataset filtered at a rigid Q-value < 0.01 and localization probability > 0.75.

In the DIA module (**Figure 1B**), PhosSight implements a spectral library pruning strategy to construct efficient, sample-specific spectral libraries (**Methods**). The workflow begins by generating a comprehensive in silico library (e.g., via DIA-NN) from a protein database, which typically contains a vast number of redundant, non-detectable precursors. To optimize this, PhosDetect is first transfer-learned (fine-tuned) using a small subset of the experimental data (e.g., a few pilot runs) to capture instrument-specific ionization patterns. The fine-tuned model then predicts the detectability of every entry in the full library. Based on these predictions, PhosSight filters the library to retain only the most “detectable” candidates (e.g., the top 50%), removing noise and reducing the search space. This filtered spectral library is then used to search the full DIA dataset, enabling faster processing and FDR control while maintaining high sensitivity.

### PhosDetect Enables Accurate Phosphorylation-Aware Detectability Prediction

To address the challenge of predicting phosphopeptide detectability, we developed PhosDetect, a deep-learning architecture that models phosphorylation-driven changes in ionization efficiency and backbone fragmentation propensities. As illustrated in supplementary Figure 1, PhosDetect employs an advanced Bidirectional Gated Recurrent Unit (BiGRU) framework that fuses sequence information with physical properties (**Methods**). The network initiates with a dual input mechanism: amino acid sequences are transformed into continuous vector representations via an embedding layer, while phosphorylation-specific physicochemical properties, including hydrophobicity, net charge, and polarity of pSer/pThr/pTyr residues are explicitly encoded. A feature fusion module integrates these two streams via a dynamic gating mechanism, allowing the model to weigh the impact of physical properties on ionization efficiency. This fused representation is processed by BiGRU layers to capture bidirectional contextual dependencies and refined by an attention mechanism to emphasize critical sequence motifs. Finally, a dense layer outputs a probability score (0-1) representing the peptide’s detectability. The model was trained on a high-quality dataset of 319,088 phosphopeptides (80/10/10 split), ensuring robust learning of phosphorylation rules (**Supplementary Figure 2**, **Methods**, **Supplementary Table 2**).

We first evaluated the generalizability of PhosDetect against state-of-the-art predictors (DeepDetect and Pfly [32]) across four standard non-phosphorylated datasets (E. coli, Human, Mouse, Yeast) sourced from the DeepDetect study. PhosDetect demonstrated superior generalization capabilities, achieving Area Under the Curve (AUC) scores of 0.89–0.96 in pre-trained modes (**Figure 2A**) and reaching 0.93–0.98 after fine-tuning (**Figure 2B**), consistently outperforming all baselines. This indicates that PhosDetect learns generalizable physicochemical determinants of peptide detectability, enabling robust performance on unmodified peptides and providing the foundation for its phosphorylation-specific predictions.

**Figure 2.**
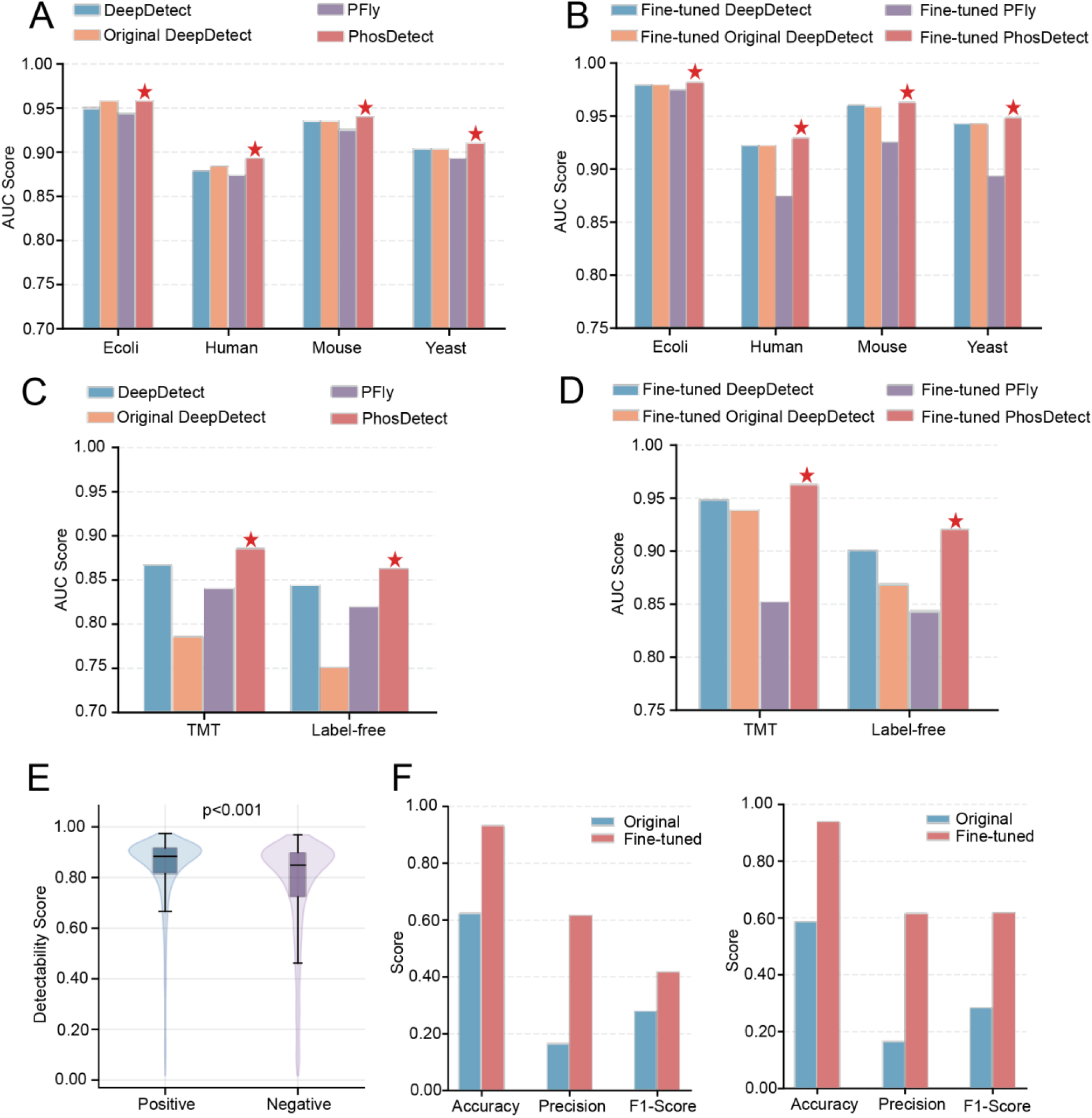
PhosDetect performance across unmodified and phosphopeptide datasets. A. Bar plot of AUC values for pre-trained models on four non-phosphorylated datasets (E. coli, human, mouse, yeast). B. Bar plot of AUC values after fine-tuning on the same non-phosphorylated datasets. C. Bar plot of AUC values for pre-trained models on three phosphoproteomic datasets (label-free, UCEC). D. Bar plot of AUC values after fine-tuning on phosphoproteomic datasets. E. Violin-box plots on a synthetic dataset, showing PhosDetect’s discriminative power between detected (positive) and alternative non-observed (negative) phosphorylation sites. F. Bar plots of accuracy, precision, and F1-score across two phosphorylated datasets, demonstrating 8–12%, 10–15%, and 9–13% improvements, respectively, after fine-tuning PhosDetect compared to baselines.

Crucially, we assessed performance on the phosphorylated peptides. Using two large-scale phosphoproteomic datasets (Label-free and TMT) sourced from the DeepRescore2 study [27], PhosDetect exhibited a decisive advantage. While generic models (DeepDetect/Pfly) showed performance drops due to their inability to model phosphate groups, PhosDetect maintained high accuracy, achieving AUCs of 0.86–0.89 (pre-trained, **Figure 2C**) and 0.92–0.96 (fine-tuned, **Figure 2D**). To further validate its discriminative power, we analyzed score distributions on a simulated test set comprising true phosphopeptides and their “non-detectable” positional isomers (**Methods**, **Supplementary Table 3**). PhosDetect showed a significant separation between positive and negative classes (p<0.001, **Figure 2E**), effectively distinguishing detected phosphopeptides from those with poor experimental detectability. In quantitative terms, the fine-tuned PhosDetect model delivered substantial gains over existing tools across the two phosphorylated datasets (**Figure 2F**). Most notably, it improved precision by 1.5 to 2.75-fold (155–275%) and F1-scores by 17–118%. This dramatic improvement in precision is particularly critical for spectral library pruning, as it minimizes the risk of discarding detectable peptides (false negatives). Collectively, these results establish PhosDetect as the dominant model for phosphopeptide detectability, providing the necessary precision to guide the subsequent DDA and DIA workflows.

### PhosSight Improves Phosphosite Localization and Identification in DDA Benchmarks

To rigorously evaluate PhosSight’s contribution to both phosphosite localization and overall identification in DDA workflows, we benchmarked eight methods on the synthetic phosphopeptide dataset [33] using MaxQuant (**Figure 3A**, **Methods**, **Supplementary Table 4**). The methods were categorized into two groups: Methods 1–4 focused on examining the impact of deep learning features on site localization accuracy without rescoring, while Methods 5–8 incorporated the Percolator rescoring module for full phosphopeptide identification. In this benchmarking framework, Method 1 (PhosphoRS) served as the conventional baseline, Method 6 represented the DeepRescore2 workflow (integrating RT and intensity), and Method 8 represented the complete PhosSight workflow (integrating RT, intensity, and detectability). The dataset comprises 96 synthetic libraries where the sequence context surrounding specific phosphorylation sites (S, T, or Y) was systematically permuted, creating a rigorous ground truth for localization validation. All methods were evaluated at the PSM level, strictly controlling the False Localization Rate (FLR) at <1% (**Methods**).

**Figure 3.**
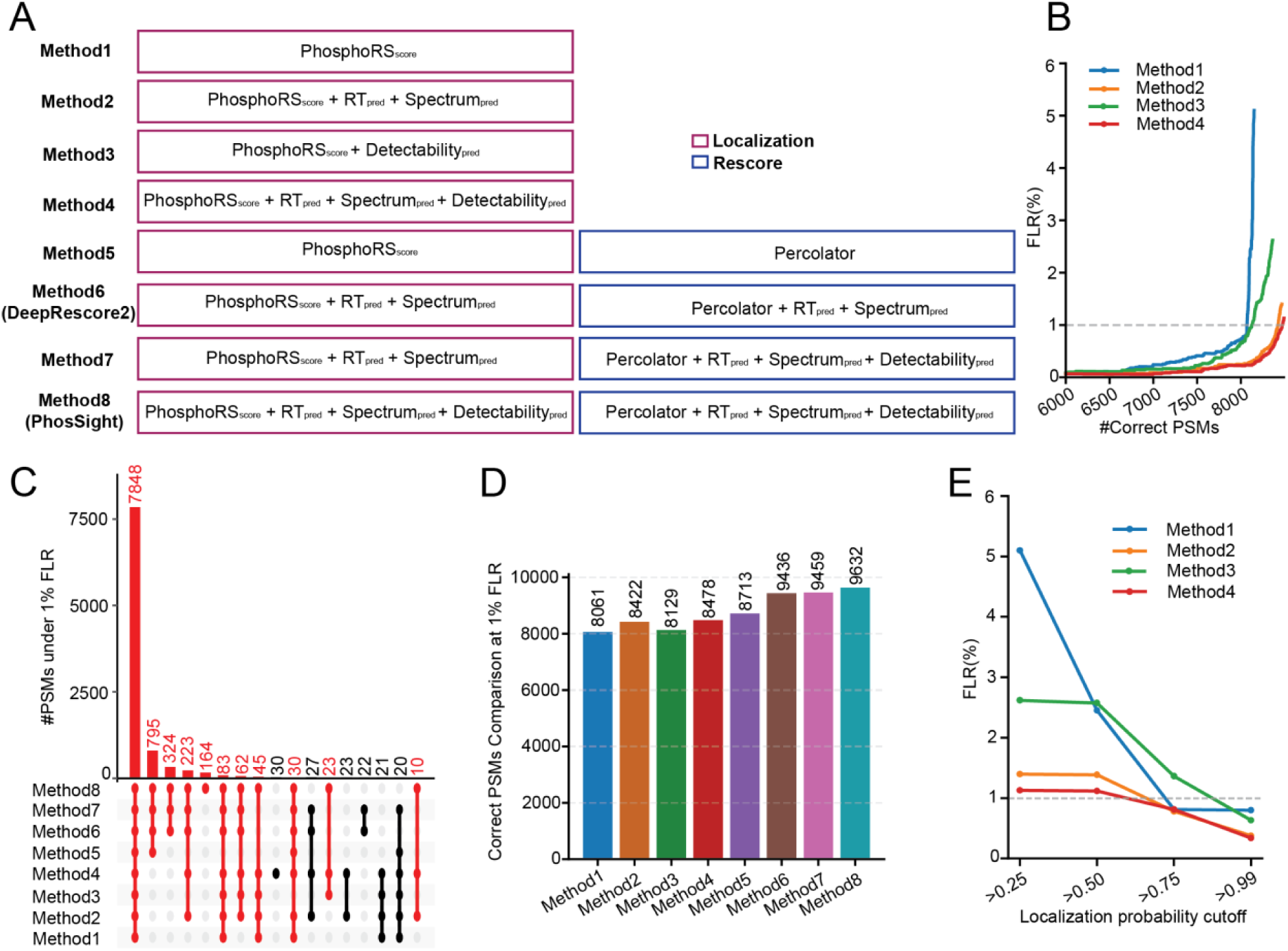
Benchmarking of deep learning-facilitated phosphopeptide localization and identification on the synthetic dataset. A. Eight Methods are benchmarked, Methods 1-4 are four site localization methods and Methods 5-8 are four phosphopeptide identification methods. B. PSM-level FLR against number of correctly localized PSMs cutoff for Methods 1-4. C. Upset plot comparing correctly localized PSMs from all eight methods at <1% FLR. The number of correctly localized PSMs by Method 8 are marked as red. D. Number of correctly localized PSMs at 1% FLR across all eight methods. E. FLR against site probability cutoff for Methods 1–4.

As shown in the FLR curves (**Figure 3B**), Method 4 (incorporating PhosDetect + AutoRT + pDeep3) consistently achieved the highest number of correct PSMs at any given FLR threshold, substantially outperforming Method 2 (AutoRT + pDeep3 only) and Method 1 (PhosphoRS baseline). This demonstrates that PhosDetect’s detectability score acts as a highly effective, independent feature that refines site localization even before rescoring. Furthermore, the upset plot analysis (**Figure 3C**) highlighted the capability of PhosSight to rescue specific, hard to identify peptides. While all eight methods shared a robust high-confidence core of 7,848 PSMs, Method 8 uniquely recovered 164 correct PSMs (highlighted in red) that were missed by all other approaches, including DeepRescore2. These unique identifications largely correspond to low-abundance or poorly fragmenting phosphopeptides where the high detectability score provided the decisive evidence for identification.

In terms of overall identification, PhosSight (Method 8) demonstrated superior performance. At a strict 1% FLR, Method 8 identified 9,632 correctly localized PSMs (**Figure 3D**), achieving the highest identification depth among all evaluated strategies. This represents a substantial 19.5% increase over the conventional baseline (Method 1: 8,061 PSMs) and a consistent improvement over the DeepRescore2 workflow (Method 7: 9,436 PSMs). The progressive increase in identifications from Method 5 to Method 8 confirms the additive value of integrating comprehensive deep learning features into the scoring engine.

Finally, evaluation across different localization probability cutoffs confirmed that Method 4 maintained the lowest FLR, particularly at the standard 0.75 threshold (**Figure 3E**). Crucially, at a localization probability cutoff of 0.75, Method 4 strictly controlled the FLR below 1% on this ground-truth dataset. Validated by this synthetic benchmark, we therefore adopted 0.75 as the standard localization probability threshold for all subsequent real-world DDA analyses. This indicates that “detectability” predictions help resolve ambiguous site assignments that both intensity and RT-based features alone cannot address. Collectively, achieving the highest total of correctly localized PSMs underscores PhosSight’s superior ability to rescue low intensity phosphopeptides in complex mixtures.

### PhosSight Accelerates DIA Search with FDR Control via Library Pruning

To evaluate the impact of PhosSight’s library pruning strategy on DIA performance, we utilized a rigorous synthetic peptide benchmark dataset [34]. This dataset mimics a complex biological background by spiking synthetic human peptides into a yeast proteome matrix at varying concentration ratios (1x to 20x). We constructed a candidate spectral library using DIA-NN, incorporating two entrapment species (E. coli and castor) to rigorously monitor false discoveries (**Methods**).

First, we assessed the fidelity of FDR control under library pruning. A primary concern with reducing library size is the potential to inflate error rates by forcing matches to incorrect targets. We calculated the entrapment false discovery proportion (FDP) across different PhosDetect filtering thresholds. As shown in Figure 4A, all filtering thresholds (from top 10% to top 90%) maintained entrapment FDPs consistently below 0.65% (nominal FDR < 1%), a level comparable to or even lower than the original unfiltered library. This confirms that PhosSight-based pruning maintains stringent specificity and does not introduce artifacts.

**Figure 4.**
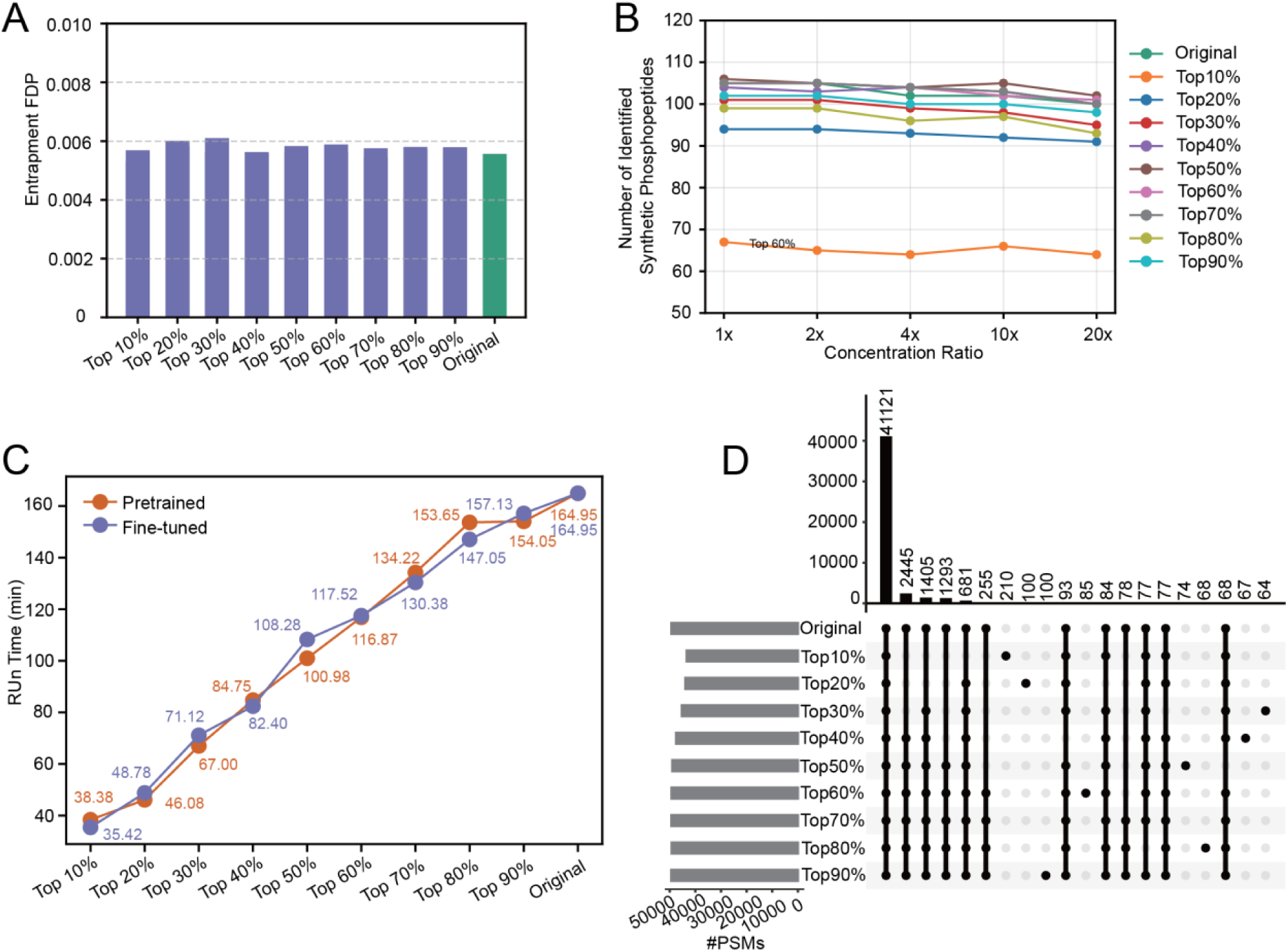
Benchmarking PhosSight in DIA data analysis using a synthetic peptide dataset. A. Effective entrapment FDP control (<1% FDR) across all detectability thresholds. B. Number of synthetic phosphopeptide identifications across concentration ratios and detectability (predicted by fine-tuned PhosDetect) filtering thresholds. C. Correlation between the spectral library size (reduced by applying different detectability thresholds) and the relative database search time. D. High overlap of peptides from searches using libraries filtered at different proportions using fine-tuned PhosDetect.

Next, we evaluated identification sensitivity. Fine-tuned PhosDetect scores were used to filter the library at various thresholds. Surprisingly, pruning the library did not simply reduce identifications. In many cases, it enhanced them by removing noise candidates. As illustrated in Figure 4B, using a library filtered to the top 90% of detectable peptides consistently yielded the highest number of identifications across all concentration ratios, outperforming the original comprehensive library. Notably, even aggressive pruning thresholds (e.g., top 60%–40%) maintained robust identification counts comparable to the original library, suggesting that a significant portion of the full library consists of “non-detectable” peptides that contribute little to signal detection.

We then quantified the efficiency gains. We observed an approximately linear dependence of database search time on the filtering threshold (**Figure 4C**). For instance, applying the top 50% threshold reduced the processing time from ∼165 minutes (Original) to ∼100 minutes, representing a ∼40% acceleration. This linear scalability implies that for larger, cohort-scale datasets, the absolute time savings would be substantial.

Finally, we examined the robustness of the method. Upset plot analysis at both the peptide (**Figure 4D**) and PSM levels (**Supplementary Figure 3**) revealed a massive overlap in identifications across different thresholds. Most peptides identified in the stricter thresholds were subsets of those found in the looser thresholds, confirming that PhosDetect consistently prioritizes the same high-quality candidates.

Integrating these metrics, we identified the top 50% detectability threshold as the optimal cutoff for general applications. It offers a dramatic reduction in search time and FDR control while maintaining identification sensitivity comparable to the full library. Consequently, this threshold was adopted for all subsequent DIA analyses in this study.

### Sensitivity Improvement in Real-World Biological DDA Datasets

To validate the robustness of PhosSight in complex biological matrices, we applied the framework to two large-scale real-world datasets: a label-free dataset from U2OS cells [35] and a TMT-labeled dataset from UCEC tumors [36]. The data were processed using four widely adopted search engines (Comet, MaxQuant, MS-GF+, and X!Tandem) under stringent criteria (1% FDR and site localization probability > 0.75, **Methods**, **Supplementary Table 5**).

At the unique phosphopeptide level (**Figure 5**), PhosSight demonstrated superior sensitivity compared to established methods. First, compared to the conventional PhosphoRS workflow (**Figure 5A**), PhosSight achieved substantial gains across all engines. For instance, in the UCEC dataset searched with Comet, PhosSight identified an additional 12,943 unique phosphopeptides while losing comparatively few (1,081) compared to the baseline. The high proportion of shared identifications (>80%) confirms concordance, while the specific gains highlight PhosSight’s ability to recover low abundance phosphopeptides that are typically discarded by standard scoring. Second, and more critically, we compared PhosSight against our previous DeepRescore2 workflow (**Figure 5B**) to isolate the specific contribution of the detectability feature. PhosSight consistently delivered additional gains, identifying between 1,000 to 2,400 more unique phosphopeptides than DeepRescore2 across the tested conditions. Specifically, in the label-free dataset with MaxQuant, PhosSight gained 1,096 peptides with negligible losses, representing a net improvement of approximately 8–15% relative to DeepRescore2. This confirms that adding “detectability” as an orthogonal dimension significantly boosts sensitivity beyond what is achievable with retention time and fragment ion intensity alone. This consistent improvement was similarly observed at the Peptide-Spectrum Match (PSM) level (**Supplementary Figure 4**). As shown in Supplementary Figure 4A, compared to PhosphoRS, PhosSight recovered a substantial number of PSMs (e.g., >9,000 gain for Comet in the Label-free dataset). Even when compared against the advanced DeepRescore2 (**Supplementary Figure 4B**), PhosSight maintained a clear advantage, recovering an additional ∼2,400 PSMs for Comet and ∼3,100 PSMs for MaxQuant in the label-free dataset.

**Figure 5.**
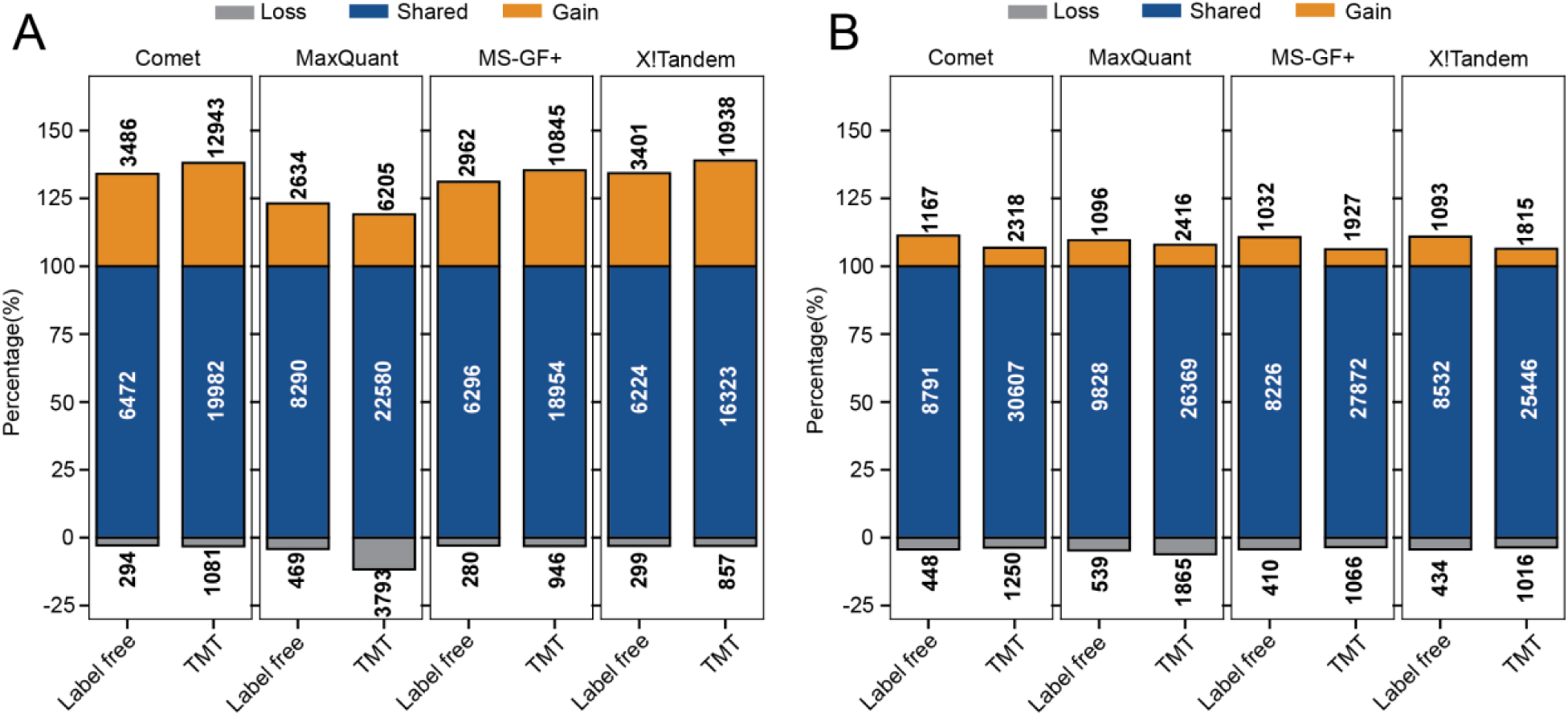
Performance comparison of PhosSight against PhosphoRS and DeepRescore2 on two real-world biological datasets label-free dataset and TMT dataset at the unique phosphopeptide level. A. The number of identified phosphopeptides from two datasets using four different search engines (Comet, MaxQuant, MS-GF+, X!Tandem) when comparing PhosSight and PhosphoRS. B. The number of identified phosphopeptides using the same four search engines when comparing PhosSight and DeepRescore2 on the same two datasets. Gain: identified by PhosSight but not by the other method. Shared: identified by both two methods. Loss: identified by DeepRescore2 but not by the other method.

Collectively, these results across four diverse search engines and two distinct quantification strategies (Label-free and TMT) establish PhosSight as a robust, engine-agnostic solution that markedly expands the observable phosphoproteome.

### Search Efficiency Improvement in Real-World Biological DIA Datasets

To validate the practical utility of PhosSight in high-throughput proteomics, we applied the library pruning strategy to a real-world human phosphoproteome dataset (A549 cells) analyzed via DIA-NN. We fine-tuned PhosDetect on a subset of the data and used it to filter the spectral library at various detectability thresholds (**Methods**).

First, we assessed computational efficiency. As hypothesized, library reduction translated directly into accelerated processing. We observed a strong linear correlation between the library size (filtering threshold) and the database search time (**Figure 6A**). For instance, applying the top 50% threshold reduced the runtime from ∼96 minutes (unfiltered) to ∼73 minutes. While stricter thresholds (e.g., top 10-20%) offered even faster speeds (down to ∼25 minutes), detailed upset plot analysis (**Supplementary Figure 5–6**) indicated that the top 50% threshold offered the optimal balance, retaining most identifications found in the comprehensive search. Consequently, the top 50% threshold was selected for in-depth comparative analysis.

**Figure 6.**
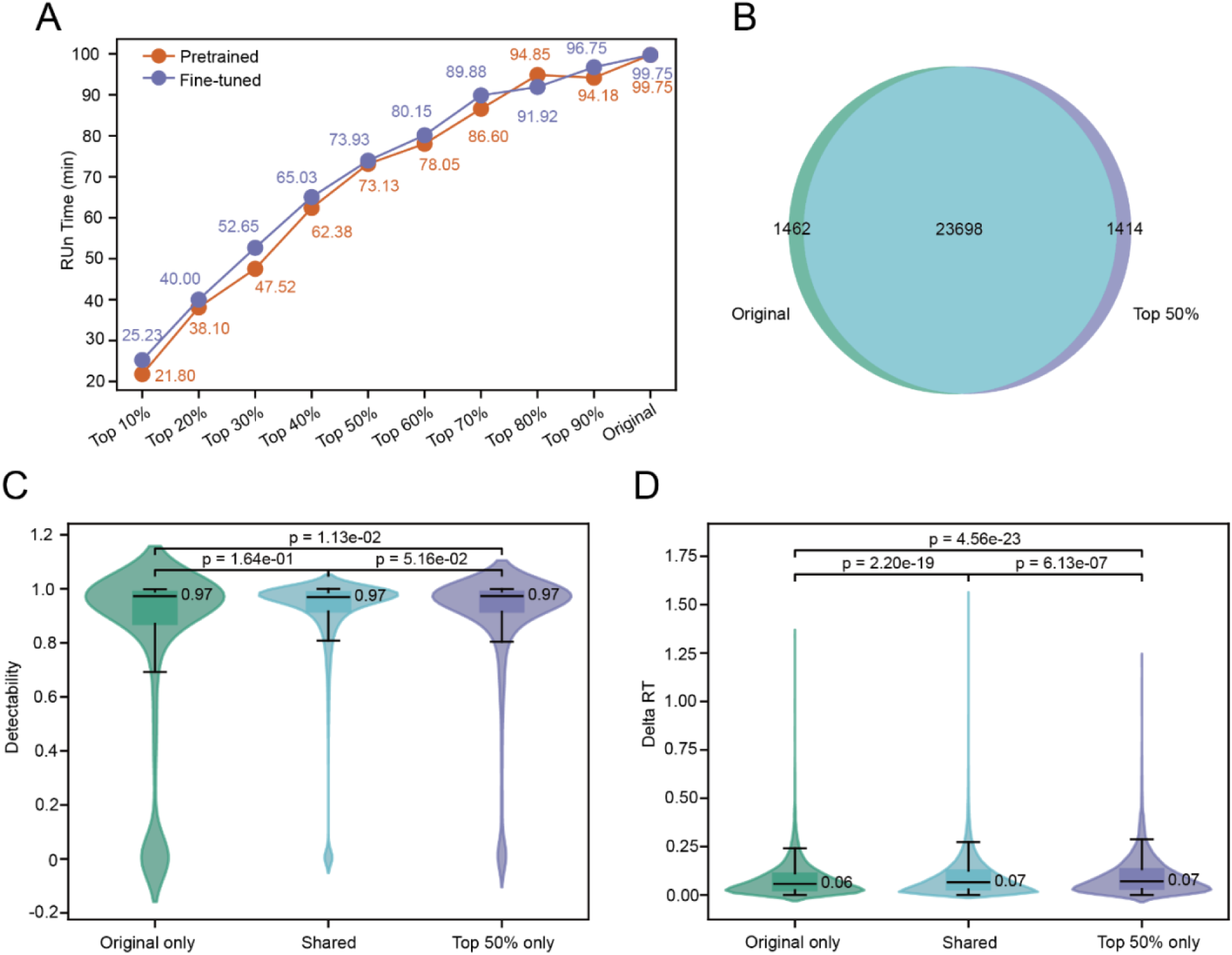
Search efficiency improvement without sensitivity loss in real-world biological DIA datasets. A. Correlation between the spectral library size (reduced by applying different detectability thresholds) and the relative database search time. B. Overlap of peptides identified from searches using the fine-tuned PhosDetect model with top 50% detectability filtering and the original unfiltered library. Detectability (C) and delta RT (D) distributions across peptide subsets from (A).

Next, we evaluated identification sensitivity and concordance. Despite removing half of the spectral library, the PhosSight-filtered search maintained an extremely high degree of overlap with the original comprehensive search. At the peptide level (**Figure 6B**), 23,680 peptides were shared between the two methods, with unique identifications constituting less than 6% of the total. This high concordance was consistently replicated at the phosphopeptide level (>13,000 shared, **Supplementary Figure 7A**) and the PSM level (**Supplementary Figure 7B–C**), confirming that PhosSight effectively captures the core identifiable phosphoproteome while discarding non-informative library entries.

Crucially, we investigated the quality of the divergent identifications. We compared the distributions of detectability scores and retention time (RT) deviations (Delta RT) for peptides found in the “Shared”, “Original only”, and “top 50% only” sets. The peptides missed by PhosSight (“Original only”) exhibited significantly lower detectability scores with a long tail extending into negative values, suggesting these were marginal identifications close to the noise floor (Figure 6C).

In sharp contrast, peptides unique to the PhosSight filtered search (“top 50% only”) possessed high median detectability scores (0.94), statistically indistinguishable from the high confidence “Shared” set. This proves that PhosSight selectively eliminates “non-detectable” peptides while preserving those with high physicochemical detection potential. We further examined the Delta RT (absolute difference between predicted and observed RT, **Figure 6D**). All three sets showed excellent RT alignment with median absolute deviations of ∼4 seconds. The “Shared” set exhibited the tightest distribution, validating the overall prediction accuracy. Importantly, the stable RT alignment in the “top 50% only” set confirms that these unique hits are likely true positives rather than random matches.

Collectively, these results demonstrate that PhosSight’s library pruning strategy (top 50%) accelerates DIA analysis by eliminating library redundancy. It preserves high-confidence identifications while preferentially selecting for peptides with higher inherent detectability, thereby enhancing search efficiency without compromising data quality.

### PhosSight Expands the Quantifiable Phosphoproteome and Uncovers Novel Clinical Associations in UCEC

To demonstrate the translational utility of PhosSight in large-scale clinical research, we applied the framework to a comprehensive UCEC cohort comprising 183 samples across 17 TMT10-plexes (**Methods**). A persistent challenge in large-scale clinical phosphoproteomics is the prevalence of missing values often caused by stochastic undersampling which fragments datasets and diminishes statistical power. We first evaluated data completeness by plotting the number of identified phosphosites against the sample coverage cutoff. As illustrated in Figure 7A, PhosSight (red curve) consistently outperformed the traditional PhosphoRS-based workflow (blue curve) across all thresholds, indicating superior data integrity and a denser signaling landscape. We subsequently defined quantifiable sites as those with less than 50% missing values within the cohort. Under this rigorous criterion, PhosSight reported 27,237 quantifiable sites, representing a 17% increase compared to the 23,232 sites identified by PhosphoRS (**Figure 7B**). Critically, this expansion was driven by the recovery of 4,630 newly quantifiable sites that were either entirely unidentified or sparsely quantified in the baseline workflow thereby significantly enlarging the search space for downstream biological discovery.

**Figure 7.**
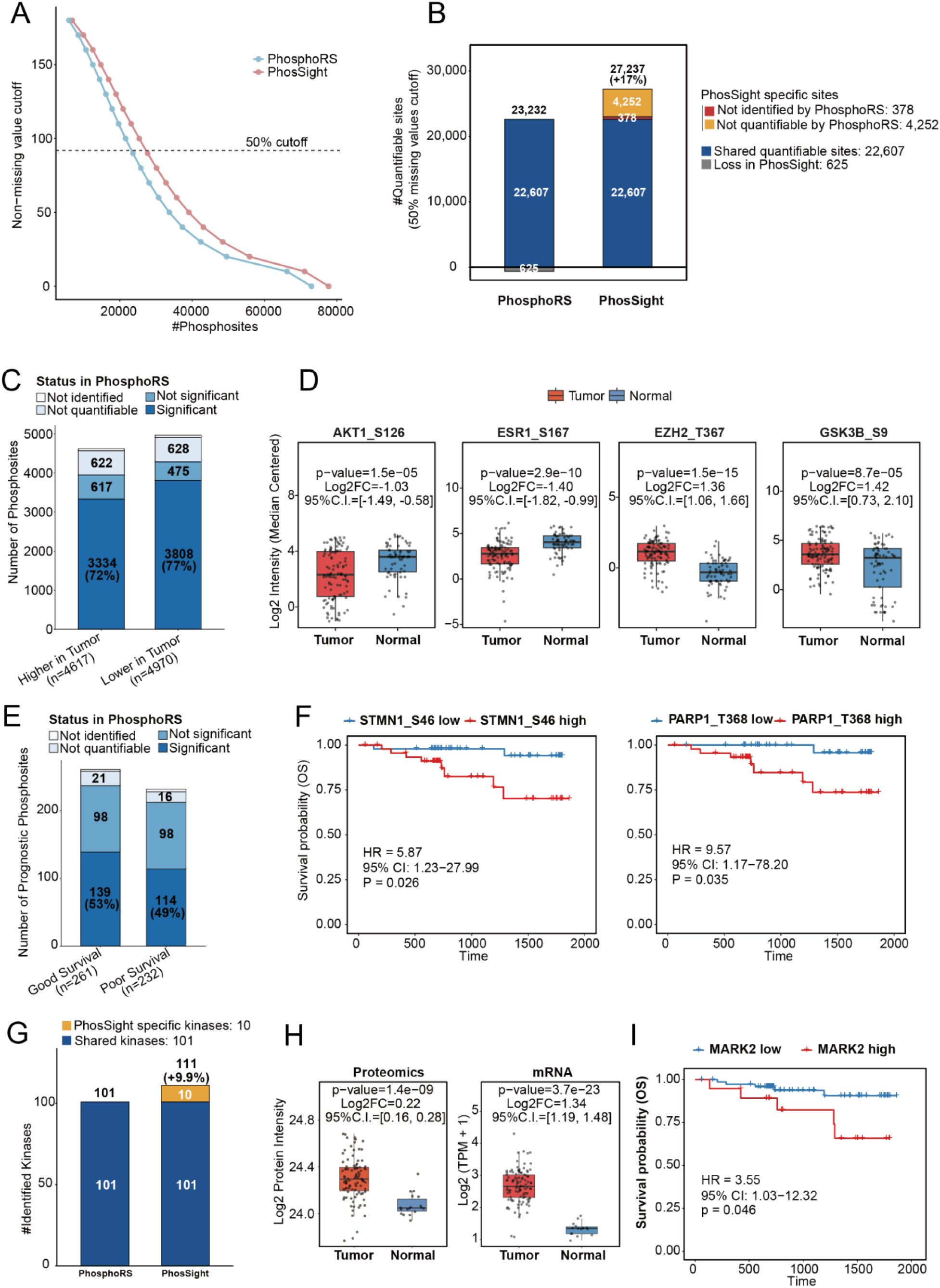
PhosSight expands the quantifiable phosphoproteome and reveals novel clinical associations in the UCEC cohort. A. Data completeness comparison showing the number of identified phosphosites as a function of the non-missing value sample cutoff. Red: PhosSight; Blue: PhosphoRS. B. Comparison of quantifiable phosphosites (defined as detection in >= 50% samples) between PhosphoRS and PhosSight. Gain: sites uniquely identified or made quantifiable by PhosSight; Shared: sites quantifiable in both workflows; Loss: sites quantifiable in PhosphoRS but not in PhosSight. C. Classification of differentially phosphorylated sites (Higher/Lower in tumor) identified by PhosSight based on their prior status in the PhosphoRS workflow, highlighting the rescue of tumor-specific signals. D. Log2-transformed intensity profiles of representative differentially phosphorylated sites (AKT1_S126, ESR1_S167, EZH2_T367, GSK3B_S9) identified as uniquely significant by PhosSight. E. Classification of PhosSight-identified prognostic phosphosites (Good/Poor survival) based on their status in the PhosphoRS workflow (not identified, not quantifiable, or not significant). F. Kaplan-Meier overall survival (OS) curves for representative prognostic sites (STMN1_S46 and PARP1_T368) uniquely captured by PhosSight. G. Comparison of inferred kinase identification coverage between PhosphoRS and PhosSight. Orange: kinases uniquely inferred by PhosSight. H. Multi-omics validation of MARK2 expression at the proteomic and transcriptomic (mRNA) levels between tumor and normal tissues. I. Clinical association of MARK2 kinase activity with overall survival. Patients were stratified into high and low activity groups based on MARK2 activity inferred from PhosSight data. (HR: Hazard Ratio; C.I.: Confidence Interval).

Leveraging this enhanced data completeness, we investigated whether PhosSight could capture clinically relevant signaling events that were previously hidden or statistically underpowered. Differential expression analysis between tumor and adjacent normal tissues revealed a robust landscape of dysregulated signaling, identifying 4,617 upregulated and 4,970 downregulated sites in tumors. A detailed status breakdown revealed that a substantial proportion of these significant hits (e.g., ∼28% of upregulated sites) were rescued from the unidentified or not significant categories of the baseline workflow (**Figure 7C**).

To validate the biological relevance of these gains, we examined specific phosphorylation sites on established cancer drivers central to UCEC pathogenesis (**Figure 7D**). Beyond general signaling trends, PhosSight’s enhanced sensitivity enabled the precise characterization of key regulatory nodes, such as AKT1_S126 and GSK3B_S9. As central effectors of the PI3K/AKT pathway, the robust quantification of these sites provides a more detailed map of the metabolic and survival rewiring occurring within the tumor. Furthermore, the framework effectively captured the differential phosphorylation of ESR1_S167, a hallmark activation site for Estrogen Receptor alpha that directly dictates hormone therapy sensitivity in endometrial cancer. Most notably, PhosSight provided high-confidence quantification of EZH2_T367 (p=1.5×10−15), a key regulatory site within the PRC2 complex that governs epigenetic silencing. By capturing such pivotal modifications, PhosSight offers deeper insights into the signaling mechanisms driving tumor progression.

The expanded depth translated directly into improved clinical stratification. We screened for phosphosites associated with patient overall survival (OS) and identified 493 prognostic sites. Notably, approximately 50% of these candidates were overlooked by the baseline workflow (**Figure 7E**), underscoring the rescue capability of our framework. This enabled the discovery of high-value prognostic markers, such as STMN1 (Stathmin 1) at S46 (Figure 7F). While previously difficult to quantify consistently, PhosSight revealed that high phosphorylation at S46, a critical regulator of microtubule dynamics was significantly associated with poor survival (HR = 5.87, p=0.026). Similarly, PARP1_T368 phosphorylation was identified as a potent predictor of poor prognosis (HR = 9.57, p=0.035). Given that PARP1 is a major target for DNA damage repair therapies, the ability to robustly quantify its phosphorylation status could provide new dimensions for precision patient stratification.

Finally, to translate site-level data into systems-level insights, we inferred upstream kinase activities. PhosSight expanded the observable kinome by 9.9%, quantifying 111 active kinases compared to 101 in the baseline (**Figure 7G**). Among the 10 kinases uniquely quantified by PhosSight, MARK2 (Microtubule Affinity Regulating Kinase 2) emerged as a standout candidate. Multi-omics integration confirmed that MARK2 is significantly upregulated at both the proteomic and transcriptomic levels in tumors (**Figure 7H**). Most importantly, survival analysis based on PhosSight-derived kinase activity identified high MARK2 activity as a significant risk factor for poor prognosis (HR = 3.55, p=0.046, **Figure 7I**). MARK2 is intimately involved in cell polarity and microtubule stability, and its identification suggests a potential role in driving aggressive tumor phenotypes in UCEC.

Collectively, these findings highlight how the enhanced coverage and sensitivity provided by PhosSight directly facilitate the discovery of actionable therapeutic targets and prognostic biomarkers that might otherwise remain obscured by technical noise.

## Discussion

Comprehensive characterization of the phosphoproteome is indispensable for decoding cellular signaling networks and identifying therapeutic targets in precision oncology. However, the inherent physicochemical properties of phosphorylated peptides, specifically their lower ionization efficiency and fragmentation suppression, have long created a trade-off between identification depth (in DDA) and computational efficiency (in DIA). In this study, we presented PhosSight, a unified framework that resolves this dichotomy by introducing peptide detectability as a critical, yet previously underutilized, dimension in phosphoproteomic analysis. Powered by PhosDetect, the deep learning model explicitly engineered to encode phosphorylation, PhosSight simultaneously boosts sensitivity in DDA workflows and accelerates library searching in DIA workflows, enabling the discovery of novel biological insights from large-scale clinical cohorts.

The cornerstone of PhosSight’s performance is the architectural innovation of PhosDetect. While generic detectability predictors like DeepDetect [29] have shown promise for unmodified peptides, our results demonstrate that they fail to generalize to the phosphoproteome due to the distinct charge distribution and hydrophobicity shifts introduced by phosphate groups. By explicitly fusing sequence embeddings with phosphorylation-specific physicochemical features, PhosDetect achieves a 2.75-fold improvement in precision compared to existing baselines.

In the context of DDA, PhosSight represents a significant evolution of our previous DeepRescore2 paradigm [27]. While DeepRescore2 successfully utilized retention time and fragment intensity, it treated all theoretical peptides as equally detectable. PhosSight corrects this assumption. This orthogonal feature proved particularly effective in identifying low-abundance regulatory sites, achieving an 8–15% increase in identification across diverse search engines. Moreover, the integration of detectability into site localization resolved ambiguities between positional isomers, where spectral evidence is often inconclusive but physicochemical stability differs.

In the context of DIA, PhosSight addresses the challenge of “search space explosion” caused by comprehensive in silico libraries. As DIA datasets grow and complexity, especially in large clinical cohorts, the computational burden and false discovery accumulation become limiting factors. Our library pruning strategy demonstrates that up to 50% of a theoretical library consists of peptides that are unlikely to be detected by current mass spectrometers. Removing these “noise” entries not only accelerates search speeds by ∼40% but also sharpens the statistical power of the analysis. Unlike random pruning, PhosSight-guided filtering preserves the core identifiable phosphoproteome, offering a practical solution for high-throughput clinical proteomics where speed and accuracy are paramount.

The translational impact of PhosSight is underscored by its application to the UCEC cohort. High missingness is a chronic issue in large-scale phosphoproteomics, often obscuring biological signals. PhosSight significantly mitigated this by expanding the quantifiable landscape. This data completeness was instrumental in uncovering the prognostic value of the kinase MARK2. While standard pipelines failed to associate MARK2 activity with survival due to insufficient substrate coverage, PhosSight’s recovery of critical substrates revealed a significant correlation between high MARK2 activity and poor prognosis. This finding highlights that technical improvements in peptide identification directly translate into the discovery of actionable clinical biomarkers that might otherwise remain hidden in the noise.

Despite these advancements, our study has limitations that warrant future investigation. First, while PhosDetect excels at predicting pSer/pThr/pTyr detectability, it is currently not trained to handle other post-translational modifications (e.g., acetylation, ubiquitination) or complex combinatorial PTMs, primarily due to the scarcity of high-quality training data for these species. As community datasets grow, extending PhosSight to a “pan-PTM” framework will be a logical next step. Second, PhosSight currently operates as a post-acquisition (DDA) or pre-search (DIA) processing tool. Integrating PhosDetect directly into instrument acquisition logic for real-time decision-making (e.g., prioritizing “detectable” precursors for fragmentation) could further revolutionize data acquisition strategies. Finally, while we demonstrated utility in TMT and Label-free settings, the framework’s adaptability to emerging acquisition methods like DIA-PASEF warrants further validation.

In conclusion, PhosSight bridges the gap between deep learning and practical phosphoproteomics. By accurately modeling the “detectability” of modified peptides, it provides a versatile, engine-agnostic solution that optimizes the trade-off between depth and speed. We anticipate that PhosSight will become an asset for the proteomics community, facilitating the comprehensive mapping of signaling networks in health and disease.

## Methods

### Construction of PhosDetect Training Dataset

The training dataset was curated from a large-scale human phosphoproteomics dataset (PRIDE ID: PXD012174) [37], processed using MaxQuant. As illustrated in Supplementary Figure 2, the data curation workflow was divided into phosphorylated and non-phosphorylated branches to ensure a balanced representation of peptide features. To ensure high data quality, stringent filtering criteria were applied. For phosphorylated peptides, only those with a false discovery rate (FDR) < 1% and a localization probability > 0.75 (“Class I”) were retained, yielding 103,945 high-confidence precursors. For non-phosphorylated peptides, a standard FDR < 1% cutoff was used, yielding 169,858 precursors.

To minimize “false negatives” arising from low protein abundance (where a peptide is detectable but missed due to instrument sensitivity limits), we implemented a protein abundance filtering strategy. For both branches, we calculated sequence coverage and peptide counts for all identified proteins and retained only those ranking in the top 50% for both metrics. From the filtered high confidence set, we selected peptides containing a single phosphorylation site (pSer, pThr, or pTyr). After filtering against the high-abundance protein list, this resulted in 79,772 positive phosphorylated samples. To generate biologically relevant negative samples (i.e., theoretically possible but experimentally undetected phosphopeptides), we utilized the PhosphoSitePlus (v2012) database [38]. Proteins from this database underwent in silico trypsin digestion (allowing up to 2 missed cleavages) to generate theoretical phosphopeptide sequences. We then performed phosphorylation site enumeration to create all possible single-phosphorylated peptides. Any theoretical phosphopeptide not identified in the experimental positive set was subsequently classified as a negative sample. This process yielded 261,590 phosphorylated negative samples.

Unmodified peptides identified in the PXD012174 dataset that passed the FDR and protein abundance filters were selected as positive samples, resulting in 161,882 non-phosphorylated positives. To generate the corresponding negative set, the UniProt Homo sapiens reference proteome was used as a background reference. Following in silico trypsin digestion (up to 2 missed cleavages), peptides that were not detected in the experimental positive set were designated as non-phospho-negative samples (n = 778,273).

The final compiled dataset comprised a total of approximately 1.28 million peptide sequences (79,772 phospho-positive, 261,590 phospho-negative, 161,882 non-phospho-positive, and 778,273 non-phospho-negative). To construct a balanced training dataset for model development, we used the number of phospho-positive samples (79,772) as a reference. An equal number of samples were randomly selected from each of the remaining three categories: phospho-negative, non-phospho-positive, and non-phospho-negative. This resulted in a final training dataset comprising approximately 319,088 peptide sequences (79,772 from each class), ensuring balanced representation across all four categories (Supplementary Table 2). This extensive corpus was used to train PhosDetect to learn sequence-specific and physicochemical determinants (hydrophobicity, charge, polarity) of peptide detectability.

### PhosDetect Model Architecture and Training

To capture the distinct ionization and fragmentation behaviors of phosphorylated peptides, we developed PhosDetect, a deep learning model based on a modified Bidirectional Gated Recurrent Unit (BiGRU) architecture. Unlike generic peptide predictors that rely solely on sequence embeddings, PhosDetect employs a dual-branch feature extraction mechanism to explicitly model the physicochemical perturbations introduced by phosphorylation. The input peptide sequence is tokenized into a 24-character vocabulary (encompassing 20 standard amino acids, 3 phosphorylated residues [pSer, pThr, pTyr], and a padding token) and processed through two parallel pathways. The first branch utilizes a learnable embedding layer to capture latent semantic dependencies within the sequence. Simultaneously, the second branch explicitly encodes three critical physicochemical properties governing peptide detectability: hydrophobicity, net charge, and polarity. To represent the chemical nature of phosphorylation, specific property values were assigned to phosphorylated residues (e.g., a net charge of -1.0 and altered polarity for pSer/pThr/pTyr) distinct from their unmodified counterparts.

A core innovation of our architecture is the implementation of a dynamic gating mechanism (Sigmoid activation) that adaptively fuses these two feature streams. This gate learns to balancethe importance of raw sequence information against explicit physicochemical constraints, generating a comprehensive feature representation. These gated features are then propagated through a BiGRU layer to capture bidirectional contextual dependencies acrossthe peptide backbone. Subsequently, a self-attention layer aggregates the recurrent outputs, allowing the model to assign higher weights to influential motifs, particularly the N-terminus and the local microenvironment of the phosphosite, which significantly impact ionization efficiency. The final weighted context vector passes through a batch normalization layer followed by a fully connected dense layer with sigmoid activation to output a probability score representing the peptide’s detectability.

The model was implemented in PyTorch and trained on a large-scale, high-quality phosphopeptide dataset constructed as described above. To ensure model robustness and prevent data leakage, the dataset was divided into training, validation, and testing sets using a stratified split strategy, maintaining consistent distributions of sequence lengths and phosphorylation states.

The training process utilized the binary cross entropy (BCE) loss function to minimize the divergence between predicted probabilities and ground-truth detectability labels. We employed the Adam optimizer with an initial learning rate of 5×10–(-4) and a weight decay of 1×10–(-5) for L2 regularization.

To achieve optimal convergence, we implemented an adaptive learning rate scheduling strategy using ReduceLROnPlateau. The learning rate was reduced by a factor of 0.7 if the validation accuracy plateaued for 8 consecutive epochs, allowing for fine-grained weight adjustments in later training stages. Furthermore, to strictly prevent overfitting, an early stopping mechanism was employed with a patience of 15 epochs. All experiments were conducted with fixed random seeds (seed=42) to ensure full reproducibility. The final model weights yielding the highest validation accuracy were retained for all downstream benchmarking and biological applications.

### Datasets

The datasets utilized in this study are categorized by acquisition mode (DDA and DIA), encompassing synthetic benchmarks for technical validation and large-scale biological cohorts for discovery.

For DDA benchmarking, we utilized a combination of synthetic and real-world datasets consistent with our previous study [27]. To evaluate phosphosite localization and identification sensitivity, we employed a synthetic phosphopeptide dataset available under the PRIDE accession PXD000138 [33]. This dataset contains 96 libraries generated from 96 seed peptides, where amino acids flanking the seed phosphorylation site (S, T, or Y) were systematically permuted to create up to 2,400 variants per library. Libraries 1 to 11 were utilized for method benchmarking, with Peptide-Spectrum Matches (PSMs) considered ground-truth positives if the sequence and localization matched the synthesized design. In addition to the synthetic data, we utilized two real-world datasets for technical benchmarking. The first was a label-free phosphoproteome dataset derived from the U2OS cell line [35], downloaded from the PRIDE database (PXD023665), from which three raw files were used to evaluate search engine performance. The second was a TMT-labeled dataset from the CPTAC Uterine Corpus Endometrial Carcinoma (UCEC) study [36], available via the Proteomic Data Commons (PDC000126). Specifically, the first TMT10-plex (comprising 12 fractions) of this cohort served as a complex biological benchmark for the PhosSight workflow. To demonstrate PhosSight’s capacity for large-scale biological discovery, we further obtained the raw mass spectrometry data for the complete UCEC cohort (PDC000126), comprising all 17 TMT10-plexes. Unlike the benchmarking subset, this full dataset was comprehensively re-processed using the PhosSight pipeline to recover low-abundance signaling events and identify prognosis-associated features across the entire patient population. All raw MS/MS data files were converted to MGF format using ProteoWizard (v3.0.19014) prior to analysis.

For DIA analysis, we employed both a synthetic multi-species benchmark and a high-quality in-house human phosphoproteome dataset. For controlled detectability benchmarking, we utilized a synthetic multi-species dataset, downloaded from the JPOST database (JPST000859) [34]. This dataset combines synthetic human phosphopeptides spiked into a yeast protein background with defined phosphorylation patterns, providing a ground truth for evaluating entrapment FDR and library pruning strategies.

To evaluate search efficiency in a realistic biological context, we generated a DIA phosphoproteome dataset from A549 cells (OEP00006796 via NODE data collection platform). Briefly, A549 cells were washed with PBS and lysed in SDS lysis buffer (1% SDS, 100 mM Tris-HCl, 10 mM TCEP, 40 mM CAA, pH 7.6). Protein extraction was performed via sonication (JY92-IIDN, Ningbo Scientz; 15% amplitude, 5 s on/off, 2 min), followed by denaturation at 95°C for 5 min and centrifugation. Proteins (1 mg) were precipitated overnight at –20°C using an acetone/ethanol/acetic acid mixture, washed, and resuspended in 100 mM NH₄HCO₃. Digestion was performed with trypsin (Promega) at a 1:50 enzyme-to-substrate ratio twice at 37°C for 16 h. Following acidification with 1% TFA and desalting using Waters Sep-Pak tC18 cartridges, phosphopeptides were enriched using a High-Select™ Fe-NTA kit (Thermo Scientific, A32992). Peptides were separated on a micro-tip column (75 μm × 200 mm, ReproSil-Pur C18-AQ, 1.9 μm) using an UltiMate 3000 nanoflow HPLC system (Thermo Fisher Scientific) with a 90-min gradient at 300 nL/min. MS analysis was conducted on a Q Exactive HFX mass spectrometer (Thermo Fisher Scientific). DIA acquisition consisted of 32 variable-width isolation windows (1 Da overlap). Full MS scans (350–1650 m/z) were acquired at 120,000 resolution (AGC 3e6, max IT 50 ms), followed by HCD-MS/MS at 30,000 resolution (NCE 27%, AGC 5e5, max IT 45 ms).

### Database searching

DDA datasets were processed using established workflows consistent with our previous study [27], utilizing multiple search engines to ensure comprehensive evaluation. For the synthetic phosphopeptide dataset, MaxQuant (v1.6.5.0) was used to search the MS/MS data against the human IPI v3.72 database concatenated with the default MaxQuant contaminant list. To maximize the recovery of candidate spectra for downstream rescoring, all default statistical filters (including PSM, protein, and site-level FDRs) were disabled. Trypsin/P was specified as the digestion enzyme with up to 4 missed cleavages allowed. Oxidation (M) and phosphorylation (S/T/Y) were set as variable modifications, with no fixed modifications applied, matching the experimental design of the synthetic library.

To assess the engine-agnostic performance of the PhosSight pipeline, the U2OS and UCEC datasets were analyzed using four independent search engines: MS-GF+ (release 2019.02.28), Comet (2018.01 rev. 4), X!Tandem (2017.2.1.2), and MaxQuant (v1.6.5.0). Searches were conducted against the UniProt human reference proteome (downloaded February 2019) concatenated with contaminants. Common parameters included: Trypsin/P digestion; Precursor mass tolerance of 20 ppm; Fragment mass tolerance of 0.02 Da. Modifications were configured as follows: Carbamidomethylation (C) was set as a fixed modification. Oxidation (M) and phosphorylation (S/T/Y) were set as variable modifications. For the TMT-labeled UCEC datasets, TMT tags (+229.1629 Da) on lysine residues and peptide N-termini were additionally defined as fixed modifications. A maximum of 2 missed cleavages was allowed for the U2OS label-free samples, whereas up to 4 missed cleavages were permitted for the UCEC TMT samples due to the complexity of the sample preparation.

DIA data processing was performed using DIA-NN (v2.2.0) with a library-free strategy (in silico library generation) augmented by PhosDetect-based pruning. For the synthetic multi-species benchmark, the search database comprised sequences of the synthetic human peptides combined with the proteomes of Saccharomyces cerevisiae (yeast), Escherichia coli, and Ricinus communis (castor). Theoretical spectra and retention times were predicted using DIA-NN’s built-in deep learning module. The search parameters were: Trypsin/P digestion with up to 2 missed cleavages; Carbamidomethylation (C) as fixed; Phosphorylation (S/T/Y) as variable (max 1 per peptide). Peptide length was restricted to 7–46 amino acids, with precursor charge states of 1+ to 4+ and an m/z range of 400–1250. Crucially, PhosDetect was integrated into the workflow to filter the generated spectral library at varying detectability thresholds prior to the final search. The search was performed with a mass accuracy of 8 ppm, MS1 accuracy of 10 ppm, a scan window of 7, and a precursor FDR of 1%.

The real-world A549 DIA dataset was searched against the human UniProt database (downloaded September 2025) using DIA-NN (v2.2.0). Similar to the synthetic workflow, spectral libraries were generated in silico with deep learning predictions. Parameters were set to: Trypsin/P digestion (max 2 missed cleavages); Fixed Carbamidomethylation (C); Variable Phosphorylation (S/T/Y, max 1 per peptide). The peptide length range was 7–46 residues, precursor charge states 1+ to 4+, and the precursor m/z window extended to 350–1650. The fragment ion m/z range was 110–1600. The in silico library was subsequently pruned using PhosDetect at the optimal detectability threshold (top 50%) to accelerate processing. Final search parameters included a mass accuracy of 10 ppm, MS1 accuracy of 4 ppm, a scan window of 10, and a precursor FDR of 1%.

### Deep Learning-Facilitated Deep Relocalization

For a candidate phosphosite i within an identified peptide sequence corresponding to spectrum S, three deep-learning derived metrics were used to adjust the PhosphoRS localization score of this candidate site i (PhosphoRSScorei).

The first deep learning-derived score is the spectrum similarity (SS) between a predicted MS/MS spectrum Sp using pDeep3 (v1.0) [30] with fine-tuning performed on experiment-specific data and corresponding experimental MS/MS spectrum Se. Based on our prior conclusion [27], we adopted Spectral Entropy [42] as the optimal metric for spectral similarity, which can be calculated as follows:

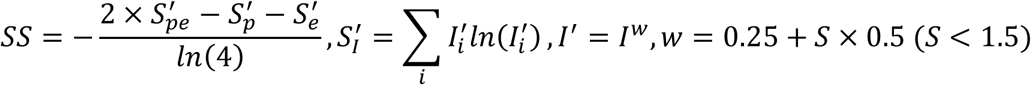

The second deep learning-derived score is the retention time difference between predicted RT (RTp) using AutoRT (v2.0) of the identified peptide sequence with phosphorylated site candidate i with fine-tuning performed on experiment-specific data as previously described [27] and the experimentally observed RT (RTe) of the corresponding spectrum. Based on our prior conclusion, we adopted RT Ratio as the optimal metric for RT difference calculation as follows:

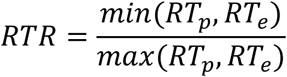

The third deep learning-derived score is the predicted detectability score (Di) using PhosDetect of the identified peptide sequence with phosphorylated site candidate i with fine-tuning performed on experiment-specific data.

For a candidate phosphosite i within an identified peptide sequence corresponding to spectrum S, the adjusted localization score is calculated by integrating the initial PhosphoRS score of this candidate site i (PhosphoRSScorei) with three deep learning-derived metrics:

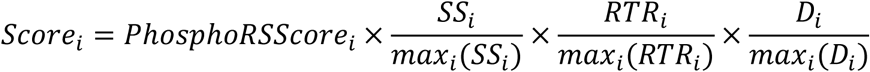

where 𝑚𝑎𝑥_𝑖_(𝑆𝑆_𝑖_), 𝑚𝑎𝑥_𝑖_(𝑅𝑇𝑅_𝑖_) and 𝑚𝑎𝑥_𝑖_(𝐷_𝑖_) are the maximum 𝑆𝑆, 𝑅𝑇𝑅 and 𝐷 for all the phosphorylated site candidates in the identified peptide sequence, respectively.

Finally, the site localization probability of phosphorylated site candidate i was calculated by transforming the localization score through the base 10 SoftMax transformation formula as previously described:

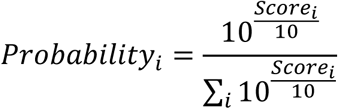

### Deep Learning-Facilitated PSM Rescoring

The localization probabilities for all potential sites are predicted and stored for each spectrum. During rescoring, the stored probabilities are used for the selection of the best phosphorylation site candidate with the largest site localization probability as the site identification of spectrum S. Following relocalization, semi-supervised rescoring was performed using Percolator (v3.4) [43] to discriminate correct identifications from random matches. For each PSM, a comprehensive feature vector (Supplementary Table 1) was constructed by concatenating three distinct sets of attributes. The first set comprised standard search engine-specific metrics, such as XCorr, DeltaCN, and Mass Error (or their equivalents depending on the search engine used). The second set included search engine-independent properties, such as precursor charge state and peptide length. Crucially, the third set integrated deep learning-derived features to capture the physicochemical validity of the assignment. Consistent with the DeepRescore2 protocol, this included the spectral similarity measured by spectral entropy distance and retention time alignment measured by the RT Ratio. Uniquely, PhosSight augmented this feature set with the peptide detectability score, representing the raw PhosDetect prediction for the localized peptide sequence. By incorporating this multi-dimensional information, the SVM model was trained to prioritize peptides that are physically “detectable” thereby rescuing low-abundance spectra. The final PSM list was filtered at a strict Q-value < 0.01.

### False localization rate (FLR) calculation for the DDA synthetic dataset

For the synthetic dataset (PXD000138), where the ground-truth phosphorylation position is known by design, FLR was calculated to benchmark localization accuracy. For each method, the analysis was restricted to phosphorylated PSMs where the peptide sequence was correctly identified. These PSMs were then ranked in descending order based on their computed site localization probability. The FLR at a specific probability threshold was calculated using Equation:

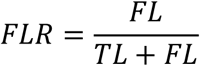

where TL represents the number of PSMs where the localized phosphosite matches the synthesized position, and FL represents the number of PSMs where the localized site differs from the ground truth, up to the given threshold.

### Entrapment false discovery proportion (FDP) calculation for the DIA synthetic dataset

To rigorously assess the fidelity of False Discovery Rate (FDR) control within the PhosSight pipeline, we conducted an entrapment analysis using the synthetic multi-species dataset (JPST000859). A composite search database was constructed by concatenating true target sequences (human synthetic peptides and yeast background proteins) with foreign entrapment sequences (proteins from Escherichia coli and Ricinus communis). Since the biological samples contained no material from the entrapment species, any identification mapping to these proteins represents a verifiable false positive. Following the DIA-NN search at a nominal 1% precursor FDR threshold, we evaluated the actual error rate by calculating the Entrapment False Discovery Proportion (FDP) among the reported identifications. The FDP was estimated using Equation:

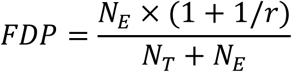

Here, 𝑁_𝑇_ is the number of reported peptides mapping to the target database (Human/Yeast). 𝑁_𝐸_ is the number of reported peptides mapping to the entrapment database (E. coli/Castor). 𝑟 represents the library size ratio of the entrapment database to the target database, used to normalize the probability of random matches.

### Phosphosite Level Quantification and Downstream Analysis for the UCEC dataset

The full UCEC cohort (PDC000126, 183 samples) was re-processed using the PhosSight DDA pipeline. To generate site-level quantification, we adopted the aggregation strategy established in our previous work (DeepRescore2) [27]. Briefly, raw TMT intensity data were extracted, and identical modified peptide sequences were aggregated by summing their abundances. The resulting phosphosite intensities were log2-transformed and median-normalized across all samples to account for technical variations. To ensure high data quality for downstream analysis, we applied a stringent filtering criterion: sites were considered quantifiable only if they possessed non-missing values in at least 50% of the cohort (i.e., detected in >= 92 samples). Data completeness was benchmarked by calculating the cumulative number of identified phosphosites at every possible sample cutoff (from 1 to 183 samples). The stability of the normalized abundance distributions across the 17 TMT plexes was assessed and visualized using PCA (FactoMineR) and density plots (ggplot2).

To identify tumor-specific signaling signatures, we performed differential expression analysis between tumor and adjacent normal tissues using the limma R package. Statistical significance was evaluated using a two-sided t-test on log 2-transformed intensities. Phosphosites exhibiting an absolute fold change |log2FC| > 1 and a Benjamini-Hochberg (BH) adjusted p-value < 0.05 were defined as significantly differentially phosphorylated. To quantify the PhosSight Advantage, we systematically compared the overlap of significant sites identified by PhosSight and the PhosphoRS baseline, focusing on the rescue of clinically relevant markers that were previously missed.

Clinical discovery was conducted by integrating quantifiable phosphosites with overall survival (OS) data. For patients with replicate tumor samples, intensities were averaged to a single value per case. Patients were stratified into high and low groups based on the median abundance of each phosphosite. The prognostic significance of each site was evaluated using univariate Cox proportional hazards regression models (survival R package). Survival probabilities were estimated via the Kaplan-Meier method, and differences between groups were compared using the log-rank test (survminer R package). Hazard Ratios (HR) and 95% Confidence Intervals (CI) were calculated, and p-values were adjusted for multiple testing using the BH procedure.

Upstream kinase activities were inferred using the Single-Sample Kinase-Substrate Enrichment Analysis (ssKSEA) approach implemented in the KSEAapp R package, with known kinase-substrate relationships sourced from the KSData repository. Phosphosite intensities were median-centered and standardized via row-wise Z-score transformation to ensure substrate variations were weighted equally regardless of their absolute abundance. For each sample, the normalized activity score for a kinase was calculated as the mean score of its quantified substrates, requiring at least three identified substrates per kinase for robust inference. To evaluate the clinical utility of these inferred activities (e.g., MARK2), we performed a systematic survival screening. Optimal cutpoints for patient stratification were determined using maximally selected rank statistics (survminer). To avoid statistical artifacts from small subgroup sizes, we enforced a constraint requiring each subgroup to represent at least 20% of the total cohort. The association between kinase activity and OS was quantified using univariate Cox regression, defining kinases with a p-value < 0.05 as significant prognostic risk factors.

## Resource availability

### Lead contact

Further information and requests for resources and reagents should be directed to and will be fulfilled by the lead contact, Xinpei Yi.

### Materials availability

This study did not generate new unique reagents.

### Code availability

The source code for the PhosSight framework, including pre-trained PhosDetect model weights, fine-tuning scripts, and comprehensive documentation, is freely available on GitHub at https://github.com/YiCITI/PhosSight. The software is distributed under the standard MIT license to facilitate reproducibility and encourage further community development.

### Data availability

The raw mass spectrometry data and associated search results utilized in this study have been deposited in standard public repositories. The large-scale human phosphoproteomics dataset used for constructing the PhosDetect training corpus is available via the PRIDE partner repository under accession number PXD012174. Datasets employed for DDA benchmarking include the synthetic phosphopeptide reference library (PXD000138) and the label-free U2OS cell line dataset (PXD023665). The CPTAC Uterine Corpus Endometrial Carcinoma (UCEC) cohort data were accessed through the Proteomic Data Commons under accession PDC000126 and PDC000199. For DIA benchmarking, the synthetic multi-species dataset is available at the JPOST repository under accession JPST000859. The A549 DIA phosphoproteomic data is available at OEP00006796 via NODE data collection platform (https://www.biosino.org/node/project/detail/OEP00006796). Processed datasets and PhosDetect predictions are accessible at https://zenodo.org/record/PhosSight_Data.

## Acknowledgement

This study was supported by the National Natural Science Foundation of China (Grants 22304111 to X.Y. and 32070668 to Y.F.). X.Y. acknowledges additional support from the Hundred Talents Program (Category B) of the Chinese Academy of Sciences, the ‘top Talent’ Program of the Shanghai Advanced Research Institute (Chinese Academy of Sciences), and the Fundamental Research Funds for the Central Universities (Grant YG2025QNA41). Furthermore, Y.F. was supported by the National Key R&D Program of China (Grants 2022YFA1004801 and 2022YFA1304603). We also acknowledge the computing resources and technical support provided by the Computing Platform (System 9) of the National Facility for Protein Science in Shanghai, Shanghai Advanced Research Institute, Chinese Academy of Sciences.

## Author contributions

X.Y. conceived the study and designed the methodology. X.Y., B.W., Z.C., and C.S. performed the formal analysis and wrote the original draft of the manuscript. The investigation was carried out by X.Y., J.Z., H.Z. and L.Lv. Supervision and funding acquisition were provided by L.Liu, Y.F., and X.Y. All authors reviewed and approved the final manuscript.

## Declaration of interests

The authors declare no competing financial interests.

## Supplemental Figures

**Supplementary Figure 1.**
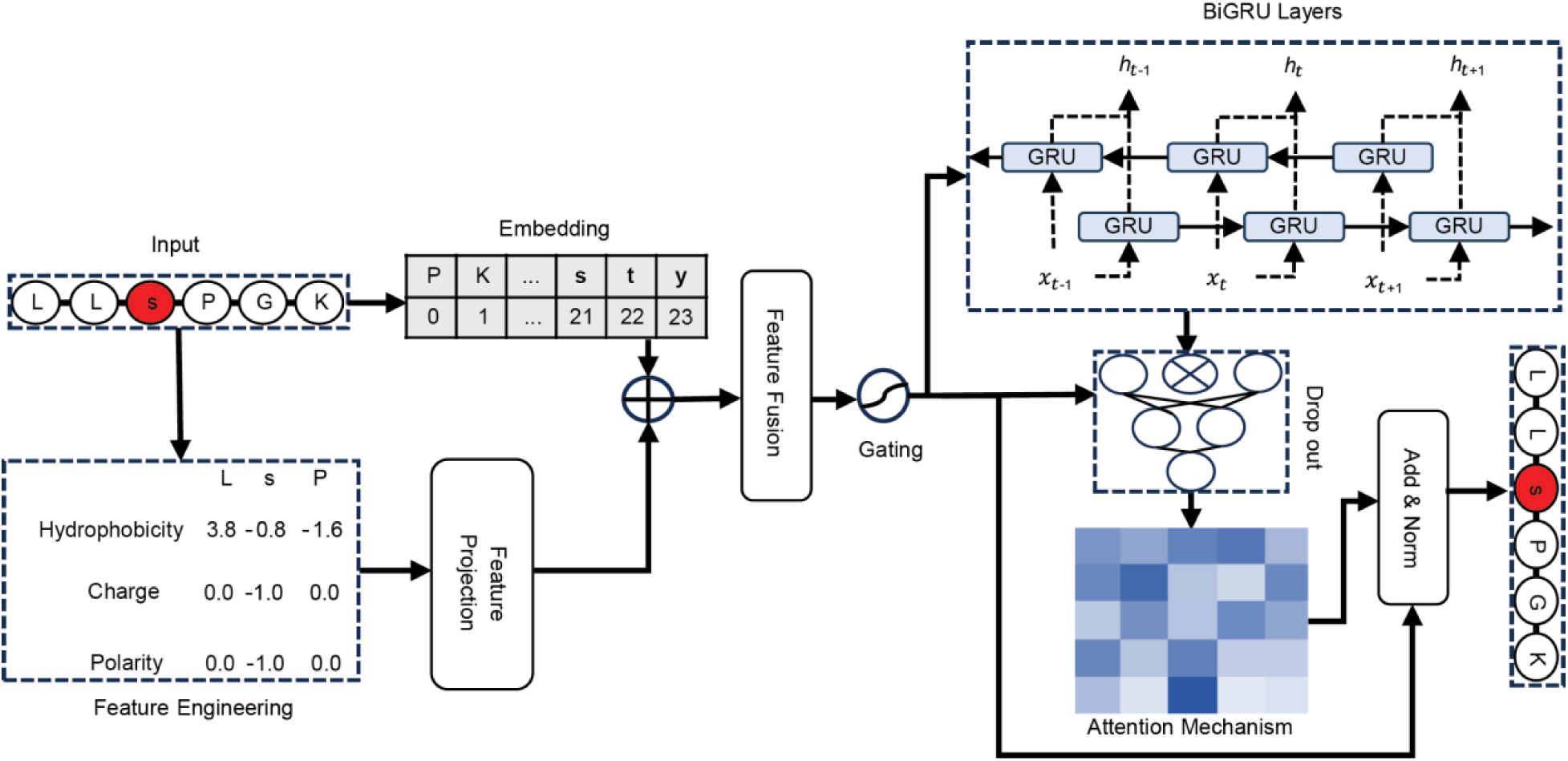
PhosDetect model architecture for pre-acquisition phosphopeptide detectability prediction.

**Supplementary Figure 2.**
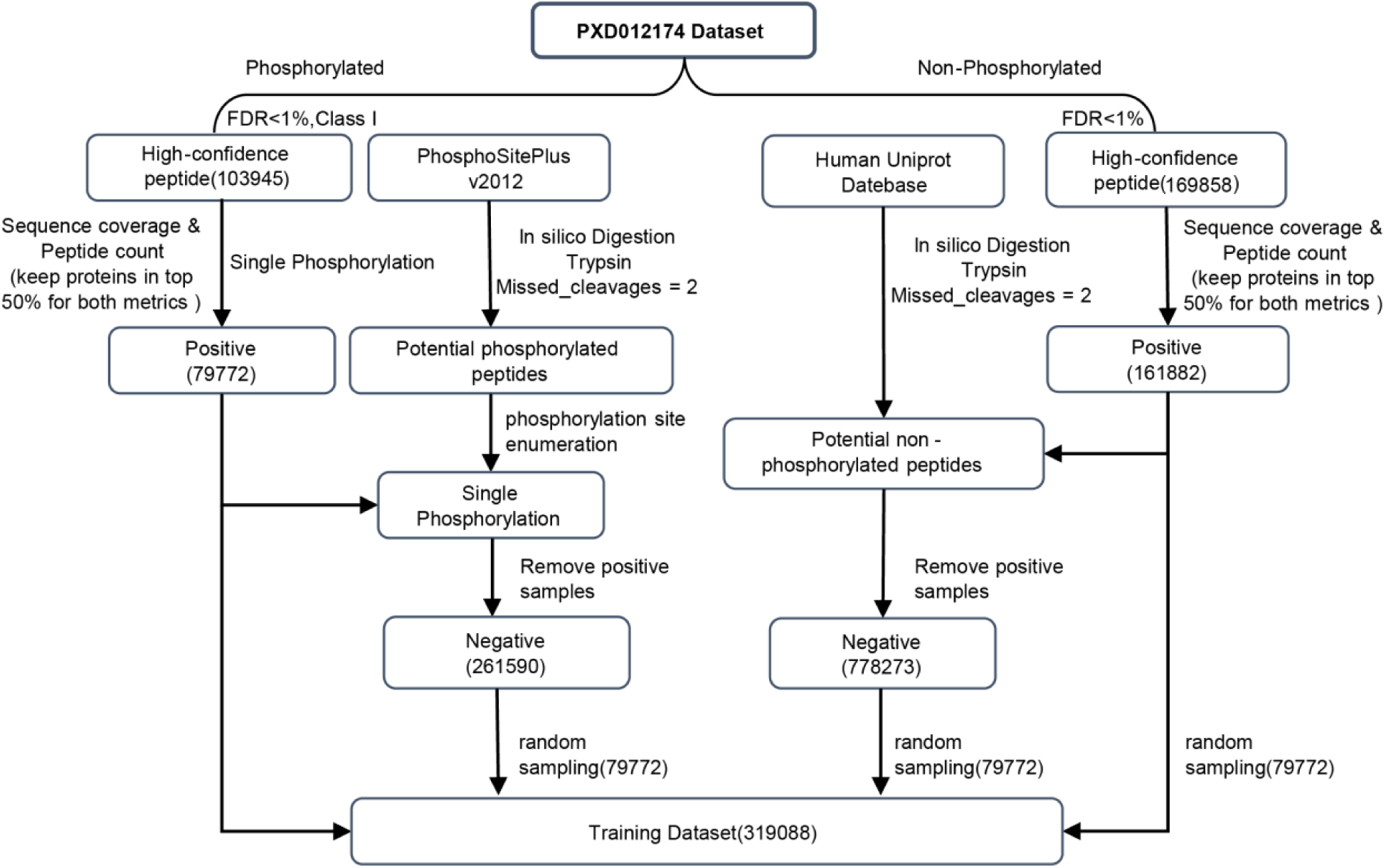
Dataset preparation workflow for PhosDetect model training.

**Supplementary Figure 3.**
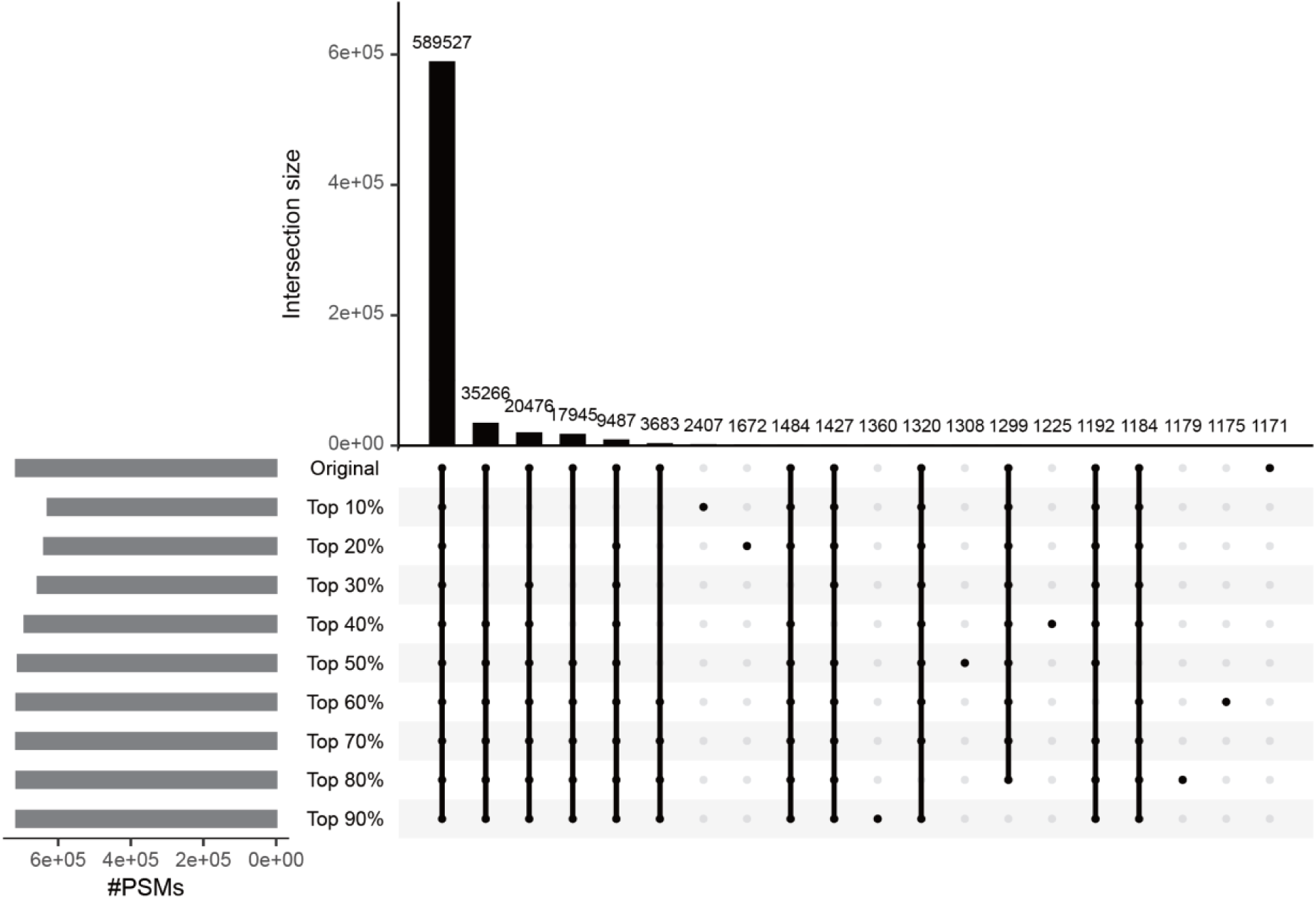
PSM overlap in DIA-NN searches using filtered spectral libraries. Upset plot displaying the overlap of PSMs across searches using libraries filtered at different proportions based on the fine-tuned PhosDetect model on dataset JPST000859.

**Supplementary Figure 4.**
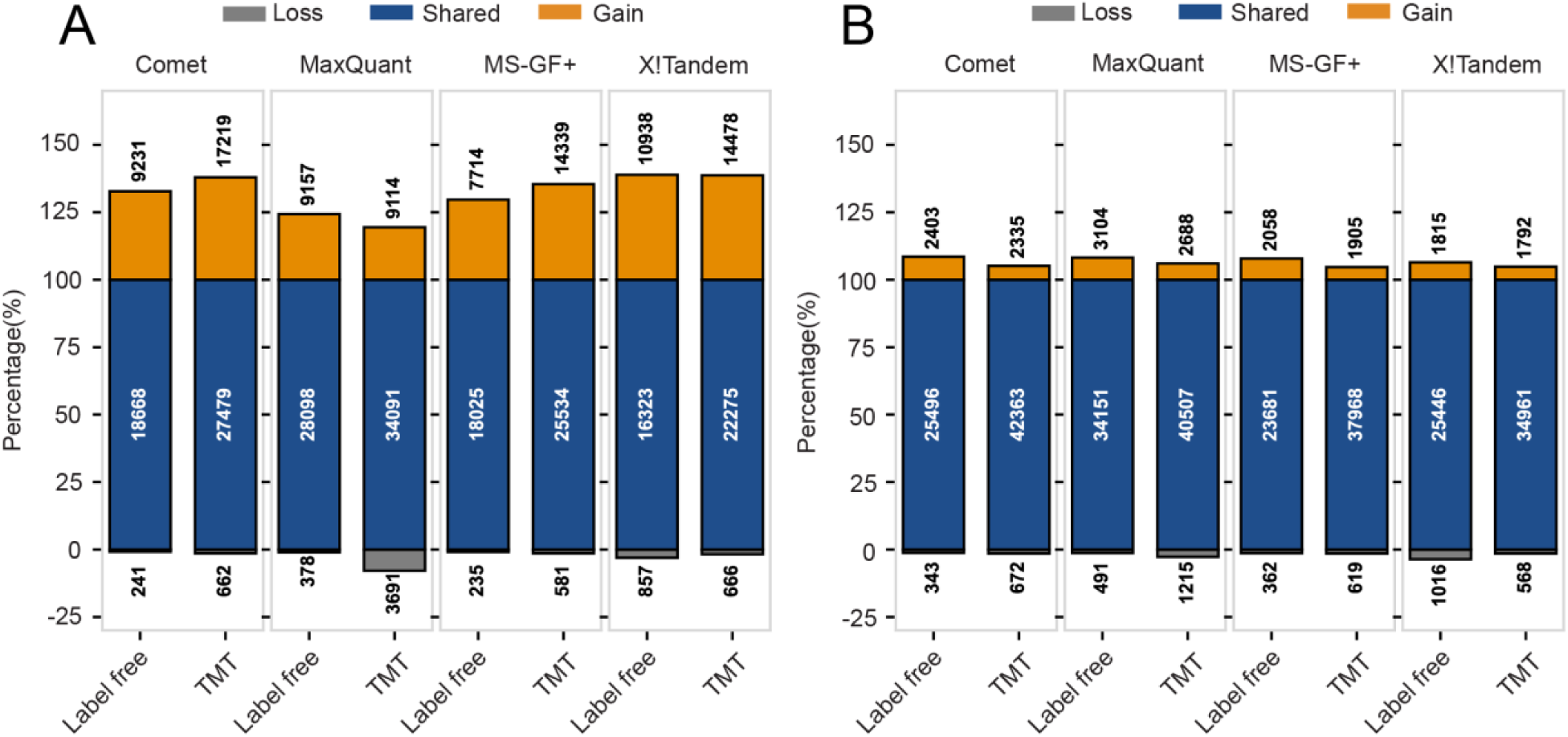
Performance comparison of PhosSight against PhosphoRS and DeepRescore2 on two real-world biological datasets label-free dataset and TMT dataset at the PSM level. (A) The number of identified PSMs from two datasets using four different search engines (Comet, MaxQuant, MS-GF+, X!Tandem) when comparing PhosSight versus PhosphoRS. (B) The number of identified PSMs using the same four search engines when comparing PhosSight versus DeepRescore2 on the same two datasets. Gain: identified by PhosSight but not by the other method. Shared: identified by both two methods. Loss: identified by DeepRescore2 but not by the other method.

**Supplementary Figure 5.**
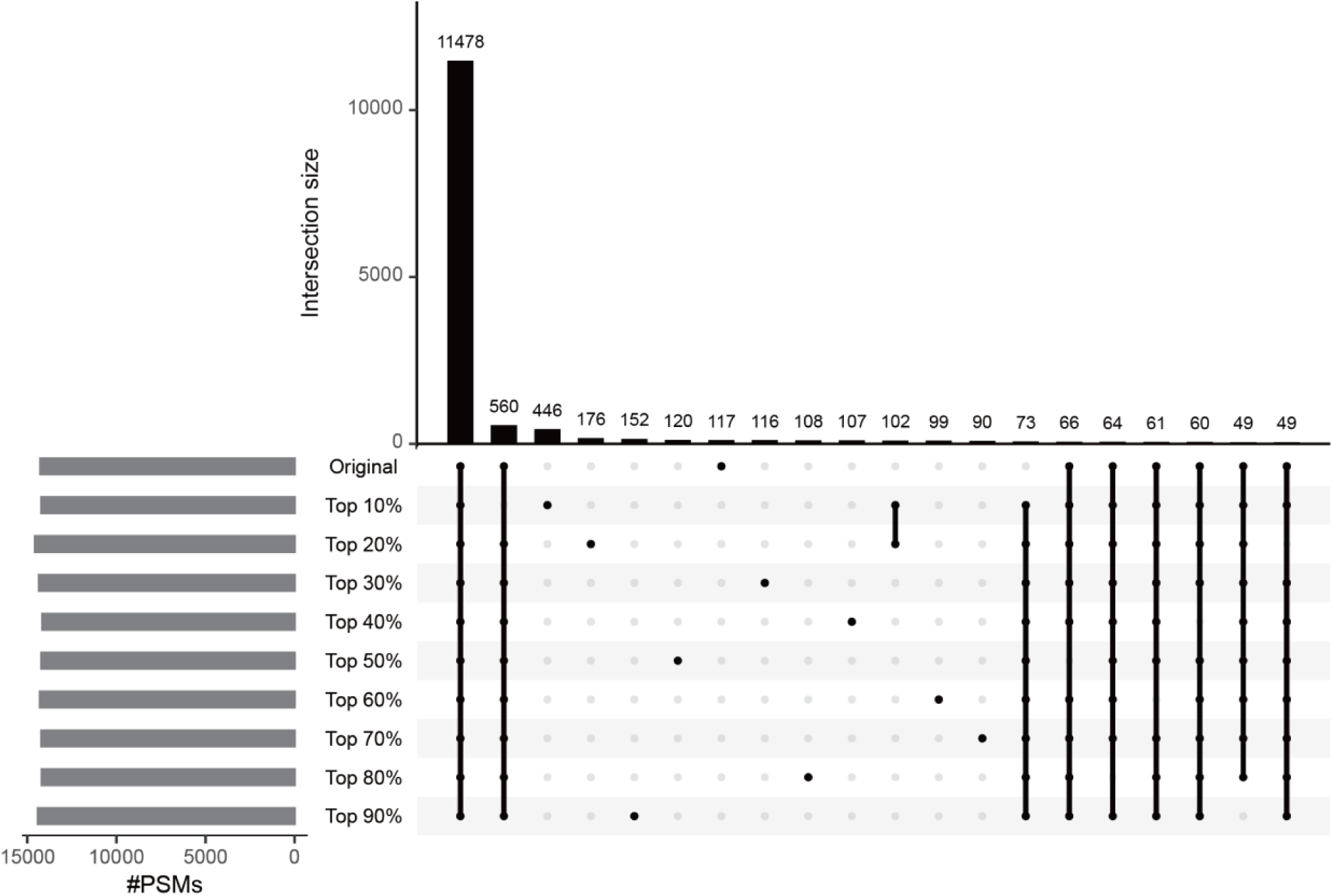
Phosphopeptide overlap across DIA-NN searches using filtered spectral libraries. Upset plot displaying the overlap of phosphopeptides identified by DIA-NN searches with spectral libraries filtered at different proportions based on the fine-tuned PhosDetect model. The analysis was conducted on A549 DIA phosphoproteomics data.

**Supplementary Figure 6.**
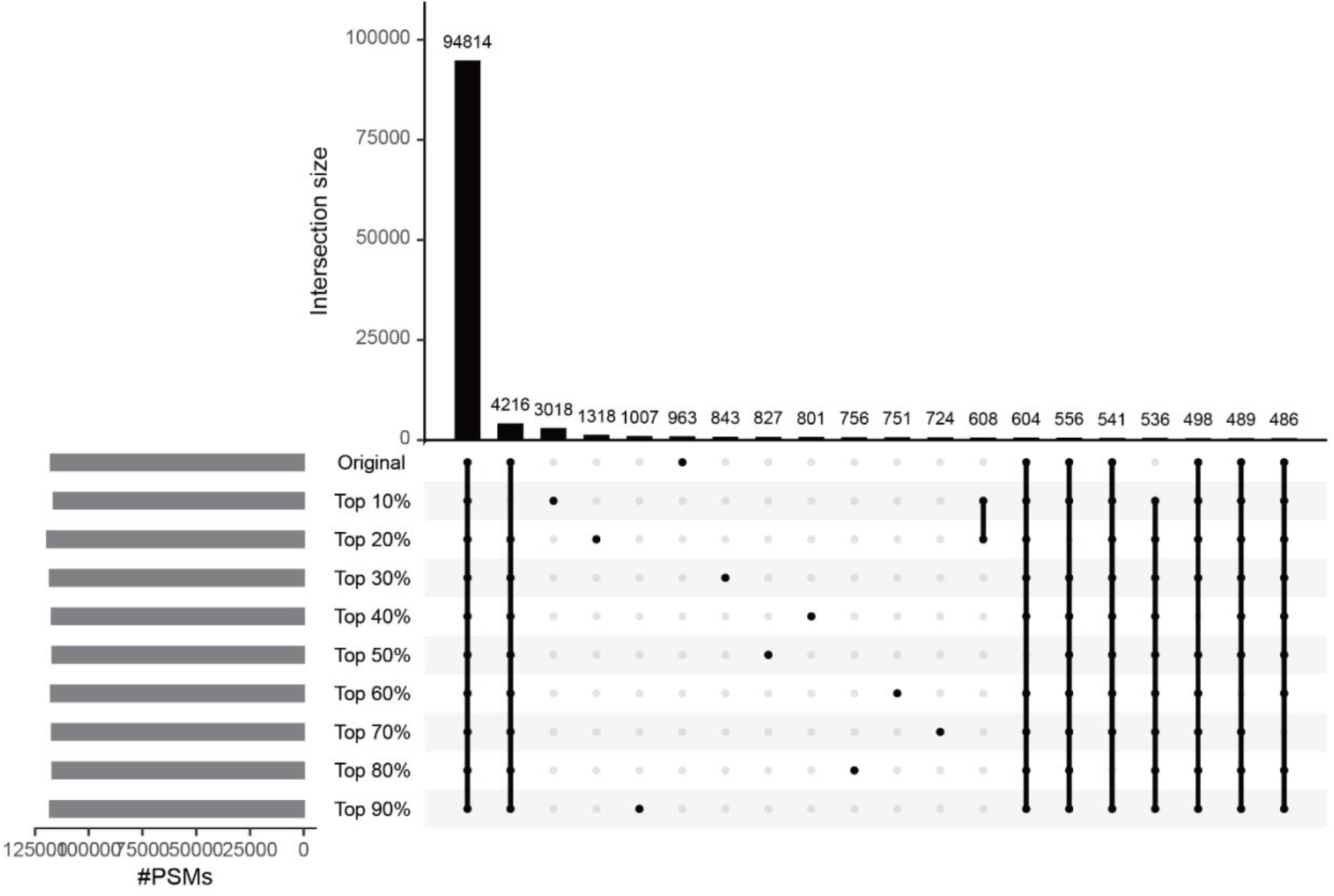
Phosphopeptide-specific PSM overlap across DIA-NN searches using filtered spectral libraries. Upset plot displaying the overlap of phosphopeptide-specific peptide-spectrum matches (PSMs) identified by DIA-NN searches with spectral libraries filtered at different proportions based on the fine-tuned PhosDetect model. The analysis was conducted on A549 DIA phosphoproteomics data.

**Supplementary Figure 7.**
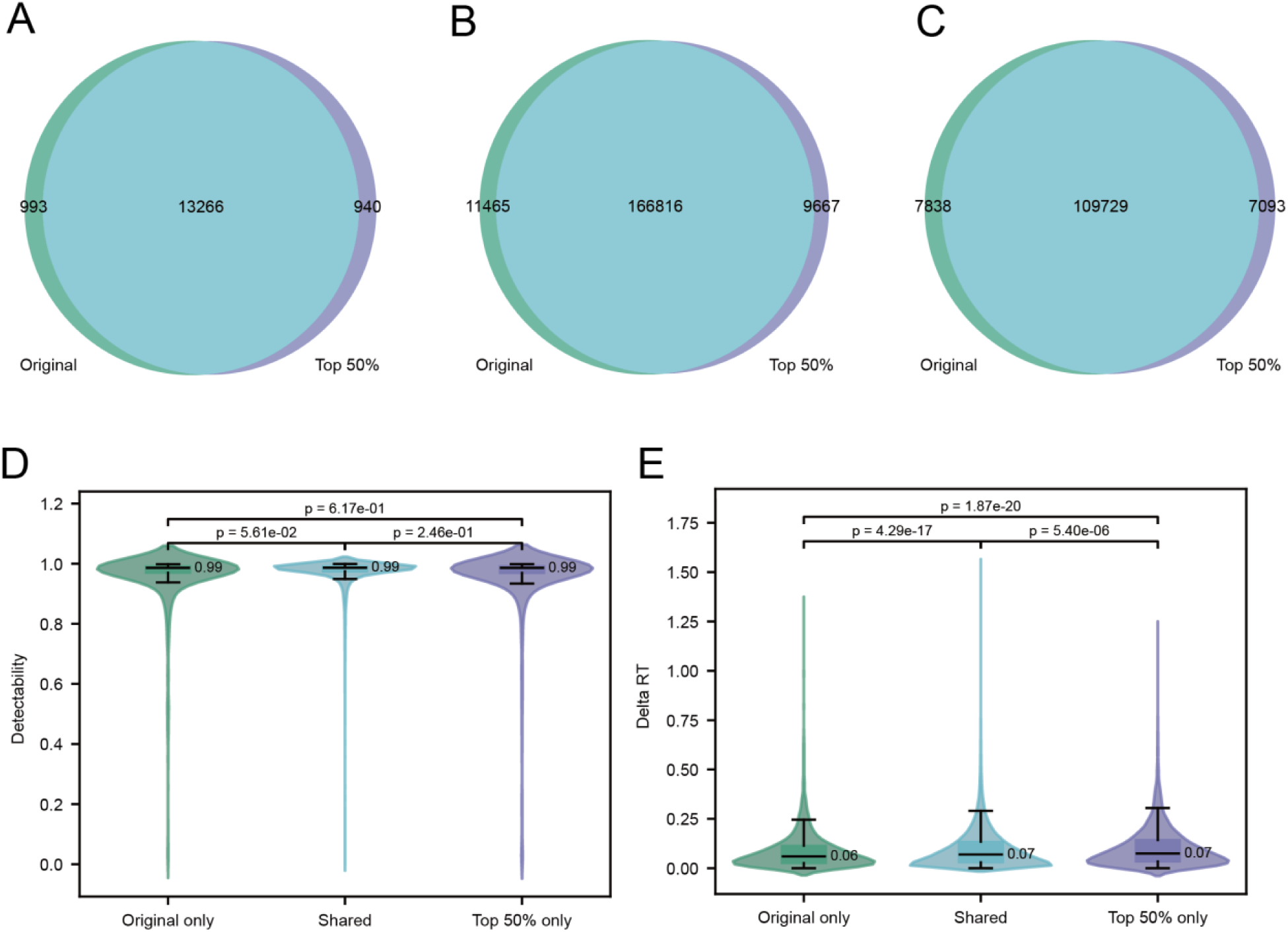
Comparative analysis of the overlap between the fine-tuned PhosDetect model (top 50% detectability) and the original unfiltered library at the phosphopeptide and PSM levels. (A) Venn diagram showing the overlap of phosphopeptide identifications. (B) Overlap of all PSMs between the two methods. (C) Overlap of phosphopeptide-specific PSMs between the two methods. Violin plots illustrating the detectability (D) and delta RT (E) distributions across the three peptide subsets (unique to fine-tuned model, unique to original library, and their overlap) defined in panel (A).

## Supplementary Tables

**Supplementary Table 1. Features used in the PhosSight workflow.**

**Supplementary Table 2. Phosphopeptide sequences and labels used for PhosDetect training.**

**Supplementary Table 3. Simulated phosphopeptide isomers used for evaluating PhosDetect discriminative performance.**

**Supplementary Table 4. Synthetic dataset results on MaxQuant search engine.**

**Supplementary Table 5. PSM identifications reported by various search engines across different identification workflows.**

## Notes

### Competing Interest Statement

The authors have declared no competing interest.

